# Whole-tissue deconvolution and scRNAseq analysis identify altered endometrial cellular compositions and functionality associated with endometriosis

**DOI:** 10.1101/2021.08.27.457966

**Authors:** Daniel Bunis, Wanxin Wang, Júlia Vallvé-Juanico, Sahar Houshdaran, Sushmita Sen, Isam Ben Soltane, Idit Kosti, Kim Chi Vo, Juan Irwin, Linda C. Giudice, Marina Sirota

## Abstract

The uterine lining (endometrium) exhibits a pro-inflammatory phenotype in women with endometriosis, resulting in pain, infertility, and poor pregnancy outcomes. The full complement of cell types contributing to this phenotype has yet to be identified, as most studies have focused on bulk tissue or select cell populations. Herein, through integrating whole-tissue deconvolution and single cell RNAseq, we comprehensively characterized immune and nonimmune cell types in endometrium of women with or without disease and their dynamic changes across the menstrual cycle. We designed metrics to evaluate specificity of deconvolution signatures that resulted in single cell identification of 13 novel signatures for immune cell subtypes in healthy endometrium. Guided by statistical metrics, we identified contributions of endometrial epithelial, endothelial, plasmacytoid dendritic cells, classical dendritic cells, monocytes, macrophages, and granulocytes to the endometrial pro-inflammatory phenotype, underscoring roles for nonimmune as well as immune cells to the dysfunctionality of this tissue.

**Teaser Sentence:** Cell type deconvolution and single cell RNAseq analysis identify altered endometrial cellular compositions in women with endometriosis

## Introduction

Human endometrium is a complex tissue that remodels during the menstrual cycle under the regulation of ovarian-derived steroid-hormones. It is characterized by phenotypic changes in diverse cell groups and changes in their relative abundance by cell proliferation and infiltration (*1*). Endometriosis is a common, steroid-hormone-dependent disorder in which endometrial-like tissue invades pelvic organs, eliciting an inflammatory response and fibrosis, resulting in chronic pelvic pain and/or infertility. The latter is due mainly to abnormal eutopic endometrium (within the uterus) that is inhospitable to embryo implantation (*2*).

Previous bulk RNAseq and microarray analyses revealed altered transcriptomic profiles in eutopic endometrium of women with versus without endometriosis (*3–5*). With disease, eutopic endometrium displays a pro-inflammatory transcriptomic feature and fails to elicit normal steroid hormone responses that are essential for endometrial transformation (*4, 6, 7*). This pro-inflammatory feature was also observed in microarray and RNAseq profiles of isolated endometrial stromal fibroblasts (eSF), mesenchymal stem cells (eMSC), and macrophages (*8, 9*). However, the full complement and abundance of cell types contributing to the pro-inflammatory feature have yet to be identified and are addressed herein.

While single cell (sc)RNAseq characterization can provide insights into phenotypes of endometrial cell populations, current costs of this technology are prohibitive for profiling samples at large scale in the context of endometrial disorders, which require sufficient sampling in both disease and controls across the diverse hormonal milieu of the menstrual cycle. Deconvoluting whole tissue level data into cell types provides a promising alternative (*10–19*) where insights such as abundance variation can be derived with high statistical power. Cell type deconvolution relies on using appropriate cell type signatures for the tissue of interest. While one strategy is to apply tissue-specific signatures derived from sorted cells or scRNAseq (*20–24*), it is limited by cell types known to the tissue, availability of signatures, batch effects from different technologies used to derive the signatures (*14*), and often does not allow for discovery of new cell types.

To leverage the advantages and overcome the limitations of these approaches, in the current study, we used both whole tissue deconvolution analysis (*25*) and scRNAseq analyses to characterize human endometrium from women with or without endometriosis. A signature compendium of 64 classical human cell types derived from diverse organs in 6 human tissue consortia were used, and a gene set enrichment-based deconvolution method was adapted (*25*). Applicability of each signature to human endometrium was evaluated by building statistical metrices using scRNAseq endometrial data obtained from women without endometriosis (*26*). In addition, to guiding data interpretation, signature evaluation prompted in-depth single cell level identification and annotation of 13 immune cell type/subtypes in healthy endometrium, including those whose identities and functions have been less well characterized and explored in endometrial biology. Herein, we present a comprehensive characterization of cellular composition of human endometrium across the menstrual cycle in women with and without endometriosis and identification of cell types with altered abundance in one or multiple menstrual cycle phases of women with disease.

## Results

### Traditional Differential Expression Analysis Identifies Immune Pathways Associated with Endometriosis Across the Menstrual Cycle

Microarray data were obtained from a public dataset (GSE51981), which was first processed and batch-corrected, followed by differential expression and pathway enrichment analyses to ensure agreement of data processing with previous literature (**Fig. 1**). **Table 1** describes the study population consisting of 105 samples across various disease stages (34 control, 24 stage I-II, 47 stage III-IV) and cycle phases (47 proliferative endometrium (PE), 24 early secretory endometrium (ESE), 34 mid-secretory endometrium (MSE)).

**Figure 1:**
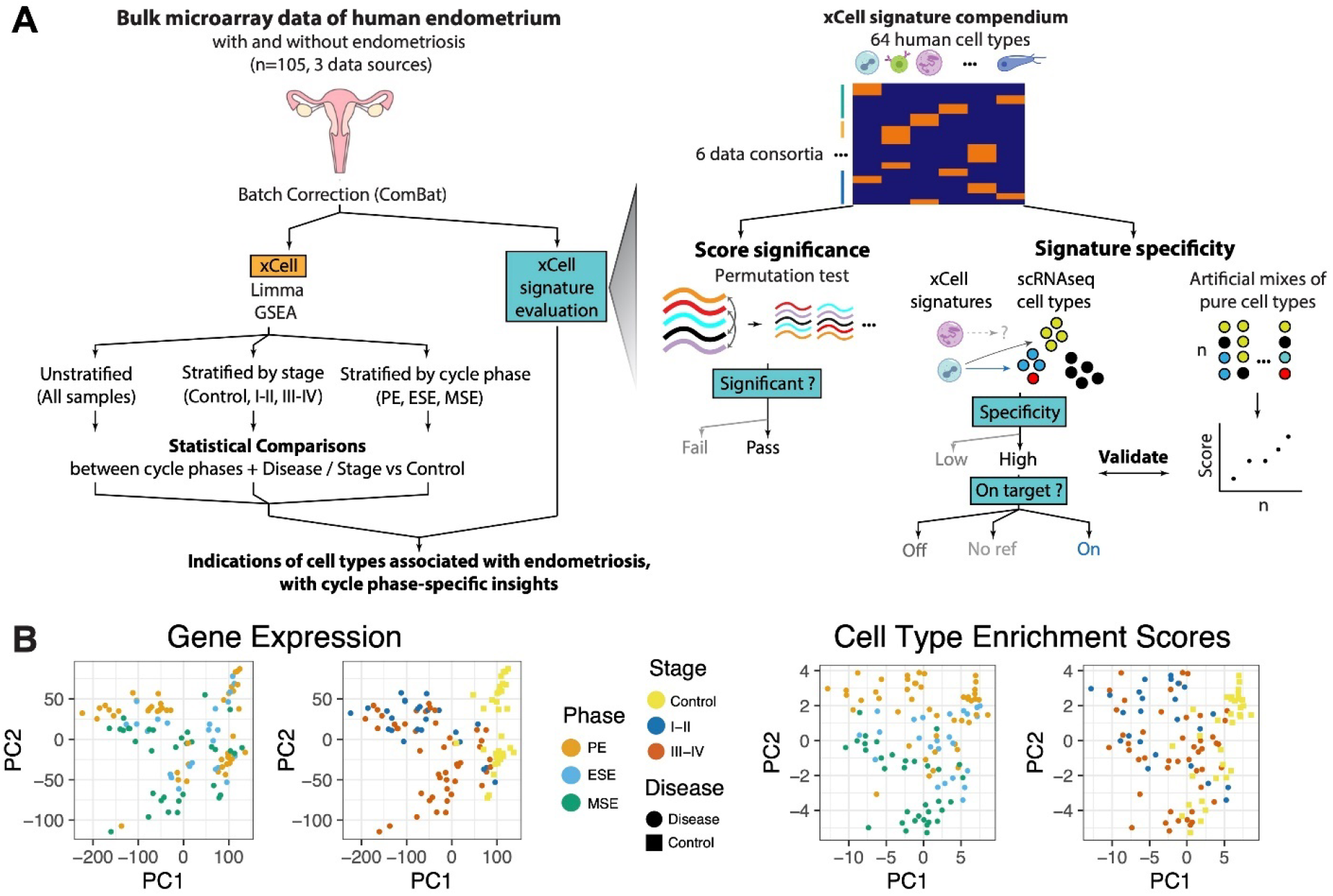
Analysis and Data Overview. **A.** Experimental overview showing how endometrial tissue transcriptome microarray data were processed and analyzed. Data were normalized (see below) and batch corrected. Then, differential gene expression (DGE) analysis was performed with log-fold change outputs run through pathway enrichment analysis. Similarly, cell type deconvolution of these bulk tissue samples was performed and validated and then analyzed for differential enrichments between sample groups. Each of these analyses was run for various stratifications (subsets) of the samples and targeting differences between distinct groupings for each stratification. These analyses were used to identify, with cycle phase-specific insights, pathways and cells associated with endometriosis. A zoom of how xCell signatures were evaluated and selected to be representative of endometrial tissue. (**Left:** Permutation test: Microarray transcriptome data were permutated at gene level to construct tissue-specific null distributions for xCell’s outputs of its 64 signatures. **Right:** Microarray transcriptome data from sorted cells were summarized and combined at different ratios into artificial mixtures. Then xCell was run on these mixtures. **Middle:** For each of the 64 xCell signatures, a two-score schematic was designed to evaluate its relationship with respect to each endometrial cell type identified in the single-cell RNAseq dataset (*26*). A specificity score (ratioNext) was agnostically quantified and each xCell signature was categorized as either targeted or non-targeted (NA: no ref) based on whether there is an endometrial cell type or subtype that the signature is targeting, and “On Target” or “Off Target” based on whether the top-ranking endometrial cell type is the signature’s intended target or not, respectively. **B.** Principal Component Analysis (PCA) of samples, after batch correction, based on (left) transcriptome data or (right) cell type enrichment scores and colored by menstrual cycle phase or disease stage.

**Table 1:** Cohort Statistics.

|  |  | Disease Severity |  |  | Lab origin |  |  | Total |
| --- | --- | --- | --- | --- | --- | --- | --- | --- |
|  |  | Control | Stages I-II | Stages III-IV | Giudice | Burney | Lessey |  |
| Cycle-Phase | PE | 20 | 10 | 17 | 31 | 13 | 3 | 47 |
|  | ESE | 6 | 6 | 12 | 14 | 10 | 0 | 24 |
|  | MSE | 8 | 8 | 18 | 19 | 11 | 4 | 34 |
| Total |  | 34 | 24 | 47 | 64 | 34 | 7 | 105 |

**Table 2.**
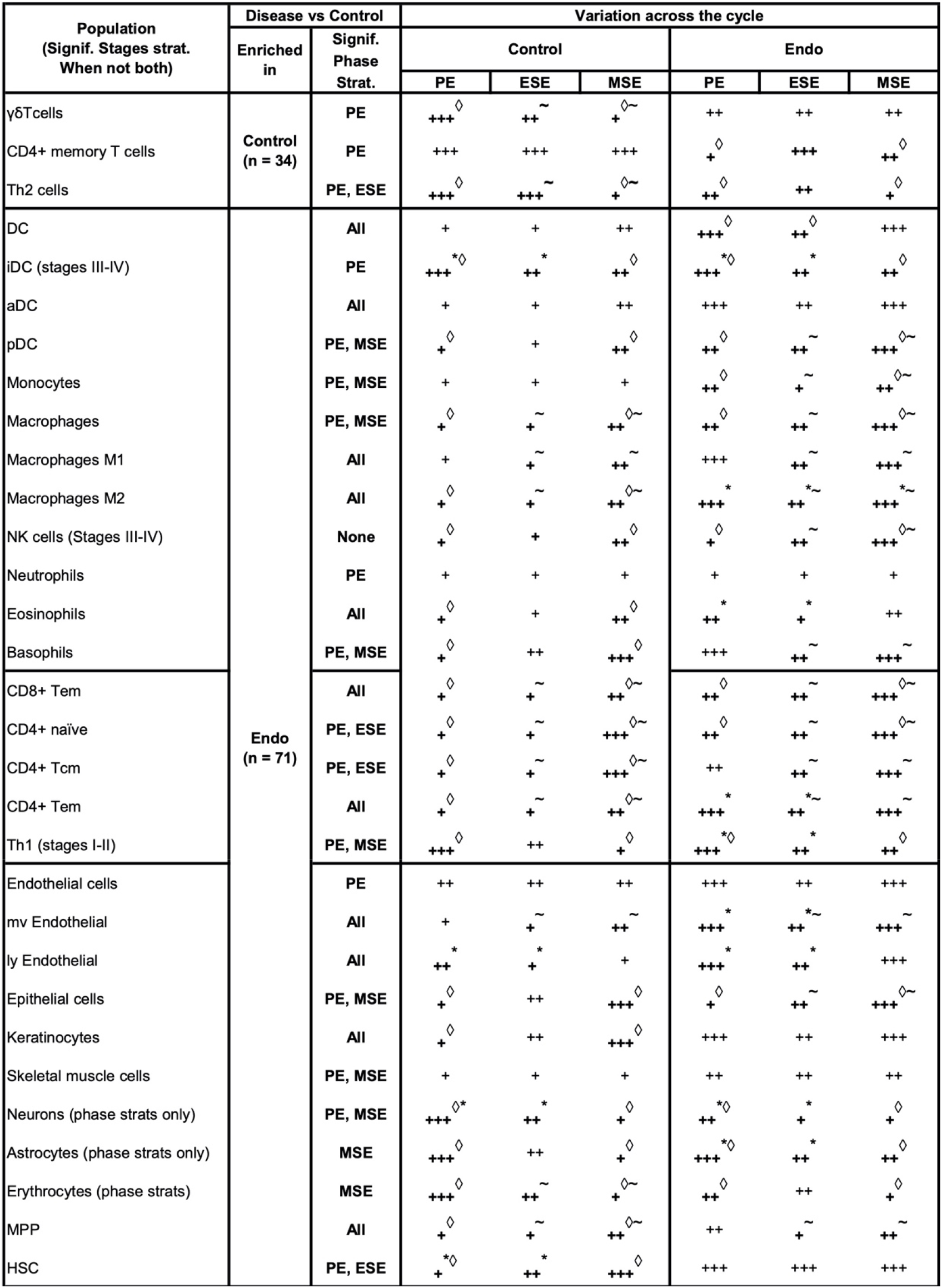
Enrichment of cell types in each condition and menstrual phase and their fluctuation throughout the menstrual cycle. Significant differences between: * = PE & ESE, ∼ = ESE & MSE, ◊ = PE & MSE Abundance of the cell type = + / ++ / +++ Unstrat. = Unstratified

Batch correction successfully mitigated laboratory-associated variations (**Fig. S1**). Based on PCA dimensionality reduction plots, samples tended to cluster by disease/stage (case) versus control and by phase (**Fig. 1B**). Heatmap clustering, focusing on genes highlighted as differentially expressed (FDR < 0.05, and log2 fold change >1) in any sample stratification, revealed strong clustering based on case versus control status (**Fig. S2A**). Phase-stratified analysis revealed overall concordance in the results (**Fig. S2B-D**). Several differentially expressed genes were identified across multiple phases, but also some phase-specific associations with endometriosis (**Fig. S2B, C**). Among significantly up-regulated genes, 79 were common to all menstrual phases such as *FOSB*, *FOS*, *JUNB*, and *EGR1*, and 182 were shared across at least two phases. In addition, there were 27 genes specific to ESE, 106 to MSE, and 428 to PE. Among the significantly down-regulated genes, 246 genes were common across all menstrual phases, including *CTSZ*, *SNTN*, *AGR3*, and *OLFM4*, and 693 genes were shared across at least two phases. In addition, there were 64 genes specific to ESE, 201 specific to MSE, and 962 specific to PE. Such discordance across phases is consistent with previous reports (*7, 27*); however, it is important to note that arbitrary differentially expressed genes (DEG) cutoffs may have amplified some of these differences as fold-fold plots revealed a high degree of correlation between phase-stratified disease versus control DEG results (**Fig. S2D**). Similarly, concordant DEG signatures were also observed when data were stratified by disease stage (**Fig. S3**).

Pathway analysis, conducted with GSEA targeting MSigDB’s Hallmark Pathways, confirmed the general concordance noted with disease versus control DEG fold-changes across stratifications and also recapitulated known biology in that many immune pathways were among those associated with disease. Interferon alpha and gamma responses, TGF-beta, and IL-2 STAT5 signaling, and complement pathways were up-regulated in endometriosis consistently across most menstrual phase and disease stage stratifications (**Fig. S4**; FDR < 0.05). TNF alpha signaling and allograft rejection, on the contrary, were significantly down-regulated in disease across all cycle phases (FDR < 0.05).

### Evaluation of applicability 64 cell type deconvolution signatures to endometrial tissue

Cell type deconvolution provides a powerful opportunity to computationally disentangle bulk transcriptomic data into individual cell types. After confirming agreement between our data processing with previous literature, we turned toward applying this technique by adapting a gene set enrichment-based deconvolution method (*25*) for use in human endometrium (**Fig.1**). The original method provided a comprehensive signature compendium for 64 classical human cell types derived from multiple organ types based on 6 human tissue consortia. To allow for discovery of new cell types and relationships between cell types, rather than using prior knowledge to select a subset of the signatures for analysis, we opted to utilize all signatures and rely on statistical metrices to infer likelihood that a cell type, of the 64, may be present in endometrium.

#### Evaluation of statistical significance of xCell output using permutation analysis

xCell produces non-zero abundance scores for all cell type signatures assessed, regardless of whether those cell types truly exist in the tissue. To overcome this, we estimated the statistical significance of enrichment scores using a permutation test (**Fig. 1A**, right). Specifically, we permuted the gene labels of the bulk data of our tissue of interest and recalculated xCell enrichment scores 1000 times to generate a dataset-specific null distribution of enrichment scores for each xCell signature. Statistical significance of scores from un-permuted data was then calculated relative to the null distribution to determine signatures for which enrichment scores were statistically significant, i.e., putatively above background. The analysis was performed separately for disease and control samples and for each cycle-phase, to ensure retention of cell types which might be abundantly present only in one condition or during a specific cycle phase (**Figs. S5 and S6**). xCell signatures passing the permutation test (ecdf_null_(median xCell score) > 90%) in at least one phase of one tissue condition were retained for further analysis and interpretation. Specifically, 50 of 64 xCell signatures passed and were deemed statistically significantly above background (**Fig. 2A**). More signatures were deemed significant in secretory phases and in the disease condition (**Fig. 2A**, **Fig. S5, S6**). In total, 22 signatures were deemed statistically significant only in disease, and only platelets were deemed above background solely in control samples. Such a finding is in line with the presumed endometrial infiltration of additional immune cell types among women with endometriosis (*28*) and during secretory phases.

**Figure 2.**
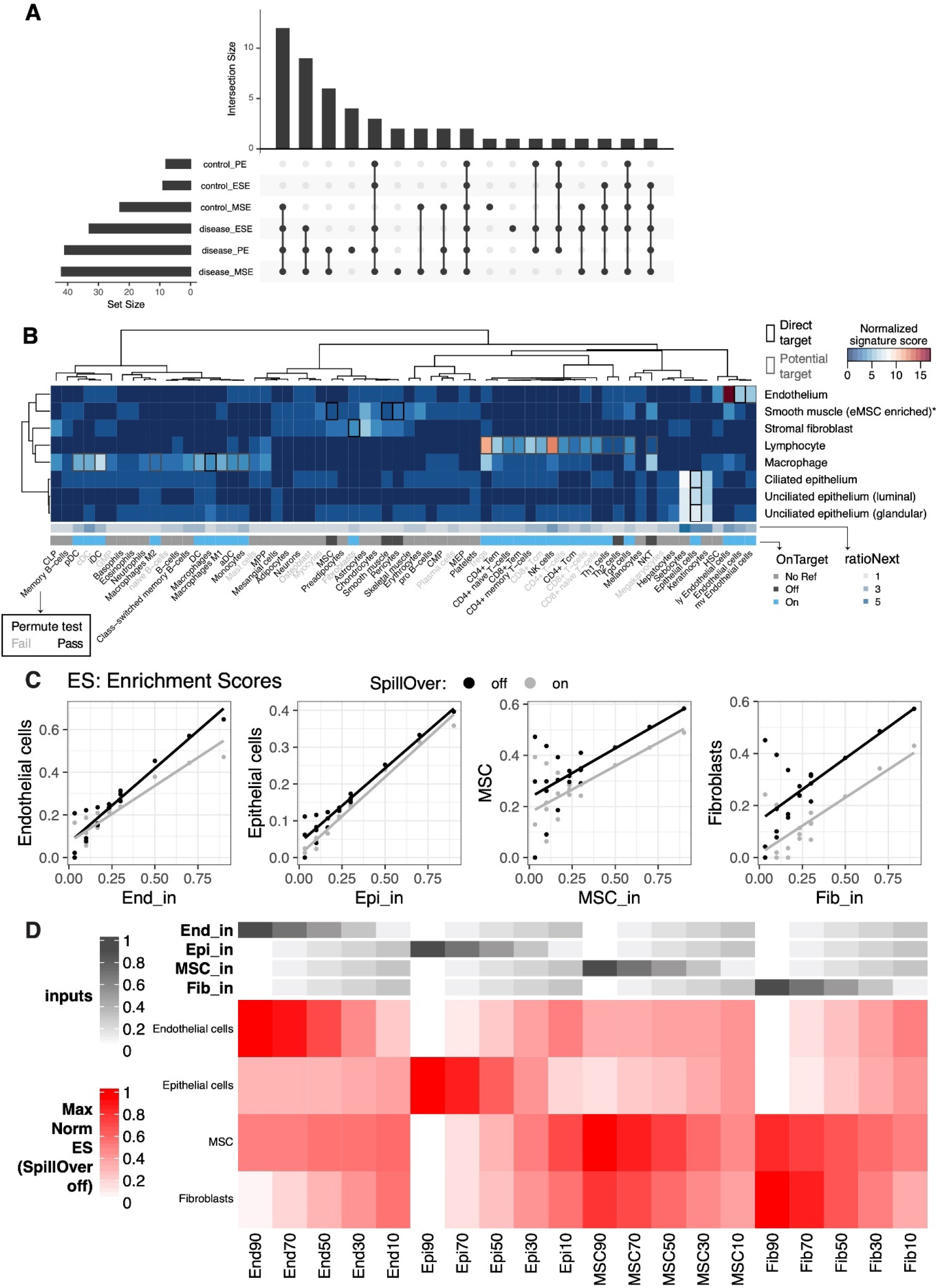
Cell Type Specific Signature Validation for Endometrial Tissue. Evaluation of xCell’s 64-cell-type signature compendium and endometrial cell types identified via scRNAseq and artificial mixtures analysis from sorted cell types. **A.** Upset plot showing patterns of conditions in which cell type signature scores were significantly higher than permuted null distributions (top), as well as the sizes of each individual set (left). **B.** Sensitivity (normalized signature score), specificity (RatioNext) and relationship (onTarget) between xCell’s 64 signatures with endometrial cell types identified at the single cell level. Shown in the heatmap are signature scores evaluated as percentage of genes in a given xCell signature that were differentially expressed between cells in an endometrial cell type compared to the remaining cells. Scores were normalized by row mediums. **C.** Scatter plots of xCell enrichment scores (y-axis) versus input percentage (x-axis), for each input cell type, with least-squares regression line overlaid, for artificial mixtures with (grey) and without (black) the SpillOver step. **D.** Heatmap of relative, normalized to the max across all mixes, enrichment scores with annotations at the top indicating input percentage of each cell type. **Abbreviations:** End= endothelial cells; Epi = epithelial cells MSC = mesenchymal stem cells; Fib = eSFs.

#### Evaluation of the specificity of cell type deconvolution signatures to human endometrial cells using single-cell RNAseq data

Even when the output of an xCell signature passes the permutation test, the associated abundance score does not necessarily reflect the behavior of its nominal target in the tissue of interest. Potential for inter-tissue transcriptomic difference of a cell type or ambiguity in naming a cell type can lead to low specificity of a signature to its nominal target. To overcome this challenge, we built two scores to evaluate xCell signatures’ specificity to (ratioNext) and relationship with (onTarget) known endometrial cell types (**Fig. 2B**, **see Methods**), using a published scRNAseq dataset of human endometrium from women without endometriosis (*26*). We plotted these two scores alongside all of our xCell outputs to help interpret the results in the context of endometrial cell types.

For xCell signatures whose direct nominal target cell type(s) were identified in the single cell dataset (**Fig. 2B**, **black boxes**), we observed high-to-moderate specificity scores with “On target” classification for all except those targeting the cell type enriched with endometrial mesenchymal stem cells (low ratioNext, “Off target”) (*29*), which we refer to, herein, as eMSCs. Specifically, we found that all candidate eMSC-targeting signatures (MSC, pericytes, and smooth muscle cells) were more differentially up-regulated in eSF than in eMSC. Results above were also validated by artificial mixtures constructed with purified endometrial cells of varying abundances, described below (**Fig. 2C, D**).

For xCell signatures without a direct nominal target cell type in endometrial tissue (onTarget=No Ref), we observed high-to-moderate specificity scores for keratinocytes, sebocytes, skeletal muscle, and hematopoietic stem cell (HSC) signatures. Given the specialized biological function of keratinocytes, sebocytes, and skeletal muscle, it is unlikely that these cell types are present in endometrial tissue. Their high ratioNext scores likely reflect the transcriptomic similarity between these cell types and endometrial cell types where they show highest specificity (e.g., endometrial epithelial cells for sebocytes and keratinocytes). On the other hand, transcriptomic similarity between HSC and endometrial endothelial cells may suggest relationship in developmental lineage.

Importantly, for many xCell signatures that do not have a direct nominal target cell type but can potentially target a subtype or a related cell type of an identified cell type (**Fig. 2B gray boxes in the heatmap**) we observed high-to-moderate specificity score and “On target” classification. Signatures that fall into this category consisted primarily of immune signatures, as well as signatures for microvascular (mv) and lymphatic (ly) endothelial cells. We reasoned that a high specificity score and a “Passed” permutation test suggest that the associated cell type/subtype likely exists in the single cell dataset but may have been concealed by more pronounced differences of major cell lineages when all cell types were included in the scRNAseq analyses. We therefore performed heterogeneity analysis on only immune cells in the endometrial dataset to explore endometrial immune cell heterogeneity and to aid further evaluation of xCell immune signatures.

#### Annotation of endometrial immune cell types at single-cell level using xCell signatures

Immune-only heterogeneity analysis revealed 13 cell type/subtypes (**Fig.3A**): 5 were from the original “Macrophage” cluster and 8 were from the original “Lymphocyte” cluster (*26*). Classical immune cell type markers allow broad annotation of these cell type/subtypes (**Fig.3B**). They alone however are not sufficient for confident cell type annotation or for measuring similarity between identified cell type/subtypes and classically defined immune cell types for scenarios described below. We therefore iterated between a signature-based scoring method (*30*) using xCell’s signatures and classical marker expression to annotate the 13 identified cell type/subtypes. Most intriguingly, we identified one cell type that stemmed from the “Macrophage” cluster but expressed classical B cell receptor component genes (e.g., *JCHAIN*, *IGKC*) at high level (**Fig.3B**). Our signature-based method revealed a distinct enrichment of plasmacytoid dendritic cell (pDC) (**Fig.3C**) and plasma cell signatures in the same cell type (**Fig. S7C**). The pDC identity of this cell type was affirmed by the expression of genes uniquely identified in pDC (*31*) such as *CLIC3* and *SCT* (**Fig.3B**) and lack of expression of classical plasma cell markers such as *CD38* and *SDC1* (*CD138*) (**Fig. S8A**). Similarly, we were able to discern among monocyte, macrophage, and classical dendritic cell (cDC) types, identify four NK cell subtypes (*NCAM1*+, *CD160*+, *CD3*+, *FCGR3A*+), one B cell type, and Tregs, whose annotation would not have been possible using classical markers alone due to marker co-expression in closely related cell groups. On the other hand, we identified one T cell subtype with high percentage of *CD8* expression (T (*CD8*+)) (**Figs. 3B & S8B**) and another T cell subtype with sparse yet unique *CD4* expression (T (*CD4*\*)) (**Fig. S8B**). xCell’s signatures for CD4+ (**Fig. S7B**) and CD8+ T (**Fig. S7E**) cells, however, were not uniquely enriched in either of these T cell subtypes. Lastly, one lymphocyte cell type distinctly segregated from the rest of lymphocytes (**Fig. 3A**) and uniquely expressed *KIT* and *IL23R* (**Fig. 3B**). We however were not able to confidently annotate it using either the signature-based method or classical markers.

**Figure 3.**
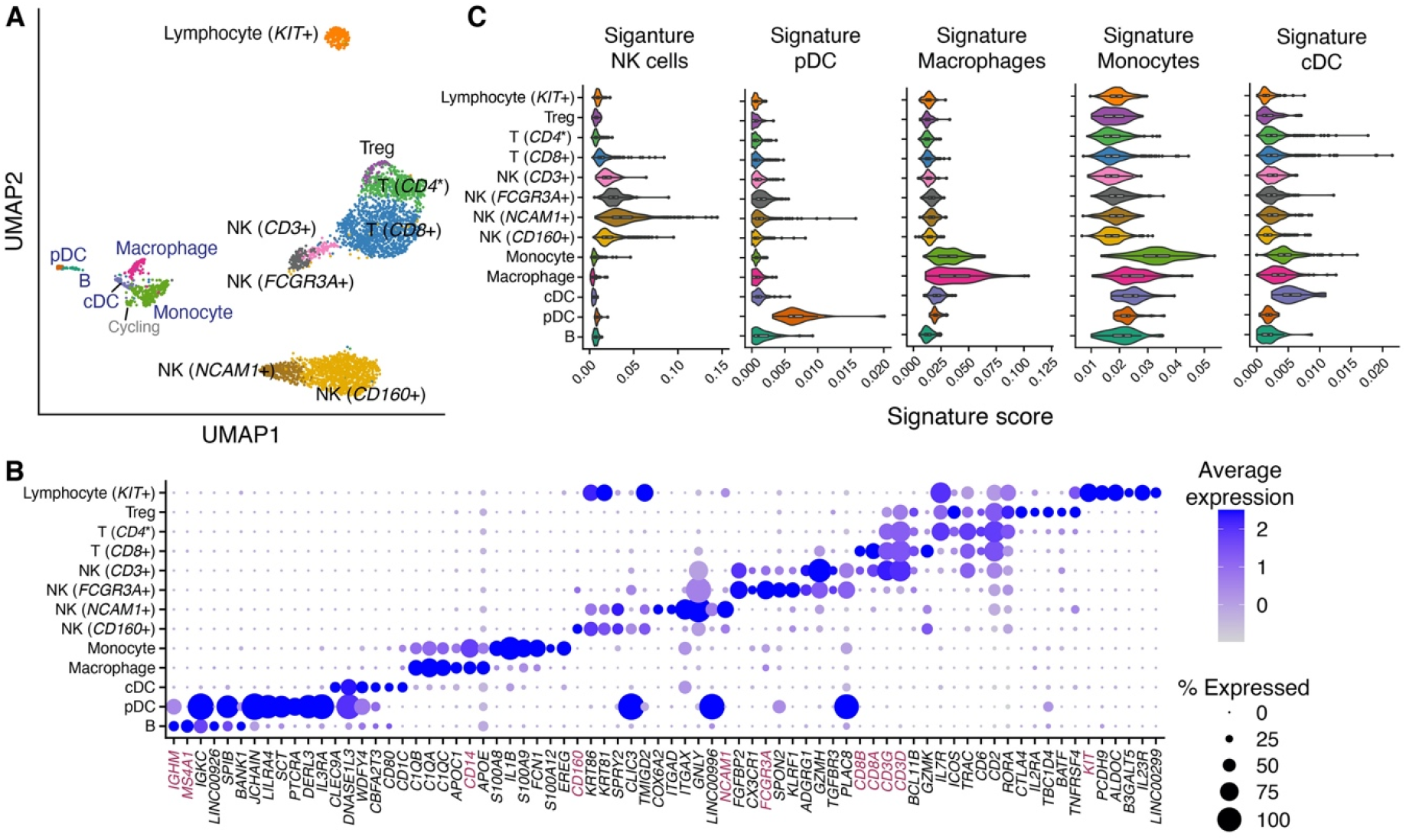
Identification and annotation of 13 immune cell type/subtypes in healthy human endometrium. **A.** Dimensional reduction and cluster identification of endometrial immune cells from women with no gynecological conditions and in natural menstrual cycles. In blue are cell type/subtypes that were from the “Macrophage” cluster and in black were from the “Lymphocyte” cluster in the original analysis. **B.** Top uniquely expressed genes for identified cell type/subtypes in (**A**). In magenta are classical cell type markers. **C.** Score distribution of selected immune signatures compiled from 6 data sources in each identified immune cell type/subtypes. (NK: natural killer cells, pDC: plasmacytoid dendritic cells, cDC: classical dendritic cells. *CD4*\*: *CD4* was uniquely but sparsely expressed in the cell subtype (**Fig. S8B**) and hence was not identified as a top uniquely expressed gene in (**B**).)

In summary, the aforementioned xCell signatures allowed confident annotation of immune cells in healthy endometrium. The unique up-regulation of their scoring in endometrial immune cell type/subtypes further confirmed their applicability in deconvoluting the tissue. Moreover, xCell’s more than 40 immune cell signatures contain not only aforementioned lineage-specifying signatures but also others that are either lineage- or function-specifying. We therefore scored all immune signatures in each of the 13 immune cell type/subtypes and plotted the result alongside deconvolution outcomes of each signature to guide interpretation (see below and **Figs. 5, S7, and S9**).

**Figure 4.**
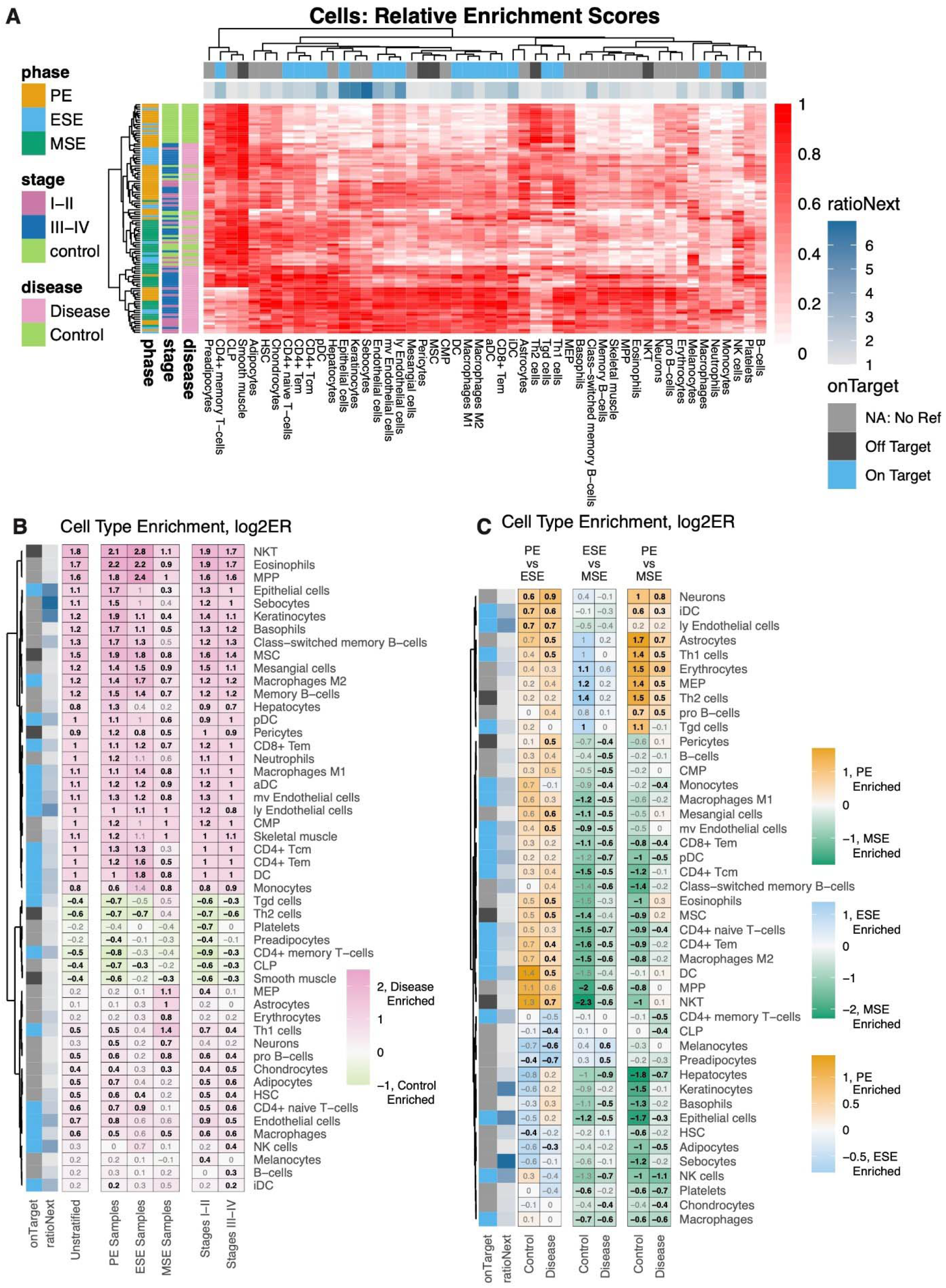
Differential analysis of Cell Type Enrichment. **A.** Heatmap showing, for all samples, relative (compared to the max per cell type) enrichment scores, of all cells determined to be differentially enriched, for any stratification analyzed. **B, C.** Cell type enrichment analysis was performed based on FDR-corrected, two-sided, Mann-Whitney U tests between (B) disease versus control for either all samples (Unstratified) or stratifications to just specific phases (PE Samples, ESE Samples, MSE Samples) or between Stages I-II (labeled as such) or Stages III-IV (labeled as such) versus control among samples from all phases or (C) between phases among case and control samples separately. Shown are heatmaps of log2 fold changes for enrichment scores where only cell types with at least one significant comparison are shown. Numbers = log2FC with black color for statistically significant enrichments and grey color for non-statistically significant enrichments.

**Figure 5.**
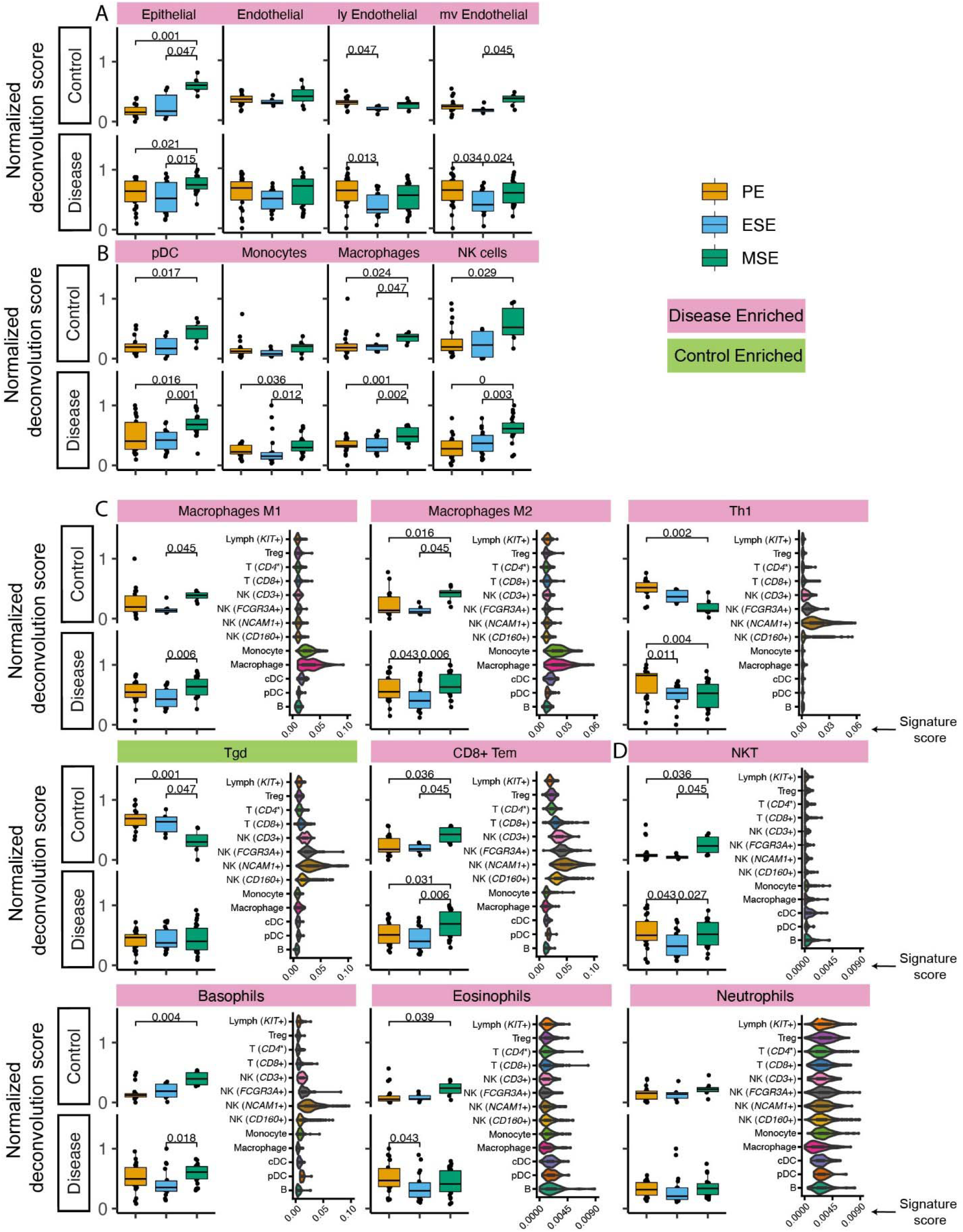
Deconvolution results and signature score distribution in single cell data of select xCell signatures. (A) with confirmed specificity to major endometrial non-immune cell types, (B) with confirmed specificity to endometrial immune cell types, (C) that are functional specifying, and (D) lack representation in healthy endometrium but show endometriosis-associated statistically significant abundance increase. Significant p-values are marked on the figure.

#### Validation of the xCell approach using artificial mixtures from sorted cells

Finally, we validated that xCell enrichment analysis could be applied to microarray-based profiles of endometrial transcriptomes by applying the approach to artificial mixtures of microarray profiles of sorted cells. Microarray expression data from four FACS-purified endometrial cell types, endothelial cells (n=11 samples), epithelial cells (n=7), mesenchymal stem cells (n=28), and eSF (n=31) (*8, 32–34*) were median summarized per gene, combined together into 20 different artificial mixtures, then analyzed with xCell (**Fig. 2C, D**). The original method has a built-in compensation step to reduce spill-over between closely related cell types. However, we observed 1) notable variations in deconvolution output in relation to which of the 64 signatures we selected as input, likely due to the use of a compensation matrix derived from *in silico* mixtures, and 2) signatures that appeared off-target in endometrial cells compared to their nominal cell type. We therefore disabled this step to ensure independence of outputs of 64 signatures. Confirming xCell’s utility, yet also the necessity of these signature assessment metrics, resulted in an overall positive correlation between input ratio and output enrichment scores (**Fig. 2C**). Yet, it also revealed an interdependence of MSC and fibroblast signatures in that enrichment scores for these signatures appeared reliant on combined input amounts of both cell types, especially with low input abundance of eMSC and eSF (**Fig. 2D**). We also confirm that removing the compensation step does not affect the overall trend in deconvolution output for these signatures (**Fig. 2C**).

### Menstrual cycle phase and endometriosis-associated changes in cellular composition of human endometrium

With metrics built for evaluating the applicability of xCell’s 64 signatures to human endometrium, we deconvoluted 105 human endometrium bulk transcriptomic profiles into the 64 cell types using the adapted xCell approach (**Fig. 1A**). Signatures that passed the permutation test were retained for downstream analysis. The data were obtained in proliferative, early secretory, and mid-secretory phases of the menstrual cycle from women with stage I-II or more severe (stage III-IV) endometriosis, as well as those without disease (control) (**Table 1**). The overall clustering of deconvoluted dataset was largely explained by disease versus control, followed by menstrual cycle phase (**Fig. 1B, right**). Cellular compositions that contributed to the changes in phase and disease were then assessed via stratified differential enrichment analysis (FDR < 0.05, no FC cutoff, **Fig. 4**) and interpreted alongside signature specificity metrices.

#### Deconvolution results for non-immune xCell signatures with confirmed specificity to cell types in human endometrium

Epithelial cells, eSF, and endothelial cells are the major non-immune cell types in human endometrium. The specificity of their associated xCell signatures to human endometrium was confirmed by our signature analysis (**Fig. 2B**), with the exception for eSF whose signature failed the permutation test. For epithelial and endothelial cells, deconvolution analysis revealed both phase and endometriosis-associated changes. Epithelial cell enrichment scores in disease were notably elevated compared to control and also varied significantly across the menstrual cycle (**Fig. 4B, 5A**). Endothelial cells were enriched in disease in comparison to control (**Fig. 4B**), with slight increase in MSE versus ESE in control (**Fig. 5A**). For both cell types, among all phases, a more significant rise in PE, compared to other phases, was observed in disease versus control (**Fig. 4B, 5A**). Disease-associated changes were also prominent in mv endothelial and ly endothelial signatures (**Fig. 5A, Fig. 4B**), both of which demonstrated high specificity to endometrial endothelial cells in our signature analysis (**Fig. 2B**). While ly endothelial signature had elevated PE scores compared to other phases in both disease and control, the PE-associated rise in mv endothelial signature was higher in disease versus control (**Fig. 4C, 5A**).

#### Deconvolution results for immune xCell signatures with confirmed specificity to cell types in human endometrium

pDC, monocytes, macrophages, and NK cells were identified in our heterogeneity analysis of single cell data from healthy endometrium (**Fig. 3**), with confirmed specificity of their associated xCell signatures (**Fig. 3C**) for deconvolution analysis. pDC enrichment scores were higher in disease over control across all cycle phases, reaching the highest enrichment score and statistically significant difference versus control in MSE (**Fig. 5B**, **Fig. 4B, C**). Similar patterns were observed for macrophage and monocyte scores, both signatures being enriched in disease across the cycle (**Fig. 5B**, **Fig. 4B, C**). In control, monocyte and macrophage enrichment scores were elevated in MSE (**Fig. 4B, C**), whereas a more statistically significant rise in MSE was observed in disease for both (**Fig. 5B**, **Fig. 4B, C**). In both control and disease, NK cell enrichment scores increased notably in MSE compared to preceding phases (**Figs. 5B & 4C**). NK scores showed slight yet statistically significant increase in stage III-IV endometriosis compared to control (**Figs. 4C & 5B**).

#### Deconvolution results for immune xCell signatures with functional applicability to cell types in human endometrium

As mentioned earlier, xCell’s comprehensive immune signatures include those that are function-specifying. Our signature-based annotation of the single cell dataset revealed that some of these signatures were uniquely enriched in cell type/subtypes that we annotated by lineage and classical markers in healthy endometrium. For example, both macrophage M1 and M2 signatures are uniquely enriched in monocytes and macrophages in healthy endometrium (**Fig. 5C**). Deconvolution results revealed an across-cycle increase in disease versus control for both signatures (**Fig. 4B**), with an elevation in MSE compared to ESE in both disease and control (**Fig. 4B**) with higher statistical significance in disease (**Fig. 5C**). Both xCell’s DC and activated DC (aDC) signatures were enriched in monocytes, macrophages, and cDC cells at single cell level, whereas immature DC (iDC) signature was elevated additionally in pDC and B cells (**Fig. S7A**). Deconvolution results revealed overall increases in DC and aDC in disease versus control, with a dip in score dynamics (significantly lower DC score in ESE vs. PE; lower aDC score in ESE vs. MSE although not statistically significant) observed in disease (**Figs. 4B & S7A**). Enrichment scores for iDC were elevated in PE compared to ESE and MSE in both disease and control (**Fig. S7A)**.

For some signatures, lineage identity of cell type/subtypes where the signatures were uniquely enriched, differed from the lineage identity of the signature. For example, Th1 signature was uniquely enriched in *NCAM1*+ and *FCGR3A*+ NK cells in healthy endometrium, Tdg signature was elevated in all NK cell types, and CD8+ Tem signature was elevated in all NK cell types and CD8+ T cells (**Fig. 5C**). Deconvolution results showed across-phase decreases in both Th1 and Tdg signatures in control but deviating behaviors in disease (**Fig. 5C**). CD8+ Tem signature had higher scores across all phases in disease compared to control and increased in MSE compared to the preceding phases in both disease and control (**Figs. 5C** & **4B, C**).

#### Deconvolution results for immune xCell signatures with low specificity to cell types in the healthy human endometrium dataset

Based on classical marker expression, we identified *CD4* expressing T cell, *CD8*+ T cell, Treg, and B cell in the healthy endometrial single cell dataset. xCell’s CD4+ or CD8+ cell type signatures that passed the permutation test did not show unique enrichment in the respective cell subtypes (**Fig. S7B**). Most of xCell’s CD4+ T cell signatures had overall elevation in lymphocytes, which explained the moderate ratioNext scores received in our signature analysis and suggests their deconvolution outcome likely reflects the collective abundance of lymphocytes.

Although xCell’s Treg and cDC signatures show moderate enrichment in Treg and cDC identified in the single cell dataset, deconvolution results of these signatures did not pass our permutation test (**Figs. 2B, S5, S6, S7A, B**).

Intriguingly, xCell signatures for B cell types generally scored higher in “Macrophage” cell type/subtypes than in “Lymphocyte” cell type/subtypes (**Fig. S7C**). This may be due to lower numbers of B cells as well as the shared antigen presenting functions between B cells and macrophages leading to joint clustering of these cell types. However, only naïve B cell signatures showed moderate elevation specific to B cells identified in the single cell dataset, yet this signature did not pass the permutation test. Enrichment scores for all B cell types were low and relatively constant in control, except for class-switched B cells, which displayed a slight increase across cycle, and for pro B-cell scores, which were elevated in PE and ESE. In disease, all B cell scores were elevated in all phases compared to control, although with a higher extent in PE compared to other phases (**Figs. 4B, C & S7C**).

#### Deconvolution results for xCell signatures lacking representation in the healthy human endometrium dataset (enriched in disease)

For several xCell signatures that passed the permutation test there were no associated cell types in the healthy single cell dataset. Thus, we have less certainty about how applicable these signatures are for potential endometrial versions of their cognate cell types. These signatures include NKT cells, neutrophils, eosinophils, basophils (**Fig. 5D**), common myeloid progenitors (CMP), and multipotent progenitors (MPP) (**Fig. S7D**), which all have relatively low enrichment scores in control (**Figs. S5** & **S6**) and consistently higher enrichment scores, across the cycle, in disease. The increase is at least two fold higher in PE for all of these cell types (**Figs. 4B, 5D & S7D**). For NKT, eosinophils, basophils, and MPPs, a statistically significant increase was also observed in control MSE compared to a proceeding phase, although xCell’s basophil signature was enriched in two NK subtypes (**Fig. 5D**).

As with our specificity analysis, keratinocyte and sebocyte enrichment scores correlated closely with epithelial cell scores (**Figs. 4A** & **S9**). HSC and endothelial signatures also correlated in the single cell dataset (**Fig. 2B**), although their enrichment score results deviated slightly in ESE (**Figs. 4A, 5A & S9**).

In both disease and control, we report a steady across-cycle decrease in erythrocyte, neuron (**Fig. S9**), and megakaryocyte-erythroid progenitor (MEP) signature enrichment scores and an overall elevation in disease (**Fig. S7D**). Increased enrichment scores were also observed for the common lymphoid progenitor (CLP) (**Fig. S7D**) signature in control and mesangial cell signature in disease (**Fig. S9**).

## Discussion

In this work, we comprehensively examined the cellular composition of human endometrium across the menstrual cycle in women with and without endometriosis, via integrated bulk tissue deconvolution and scRNAseq analysis. Our approach was uniquely designed such that we leveraged a large sample size of bulk data, a comprehensive signature compendium for 64 classical human cell types based on 6 human tissue consortia, a GSEA-based deconvolution method, and a high resolution of single cell RNAseq data - mitigating limitations inherent in each factor. Importantly, while benefitting from the comprehensiveness of the 64 signatures, we designed statistical metrics to evaluate the applicability of each to human endometrium to ensure statistical significance and guide interpretation. With this approach, we identified cell types with altered enrichment in one or multiple menstrual cycle phases of women with endometriosis versus controls without disease. Also, novel transcriptomic-level signatures for 13 immune cell type/subtypes in healthy endometrium, not heretofore reported in endometriosis, including pDC and monocytes, were identified. The positive enrichment of these transcriptomic signatures might indicate presence in endometrium of previously unidentified cell types, or even phenotypes among known cell types, that had not been previously investigated (see discussion below for NK, and T cell subtypes).

### Contributions of Non-immune cells to Endometriosis

Our signature evaluation confirmed the specificity of xCell’s signatures to most major non-immune endometrial cell types, including epithelial cells, and endothelial cells, but not fibroblasts. We observed increased enrichment scores in PE with endometriosis for epithelial cells and endothelial cells. This is consistent with observations of increased endothelial proliferation in women with endometriosis and menorrhagia versus controls (*35–37*) has been observed.

eMSC is an endometrial cell type that exhibits mesenchymal stem cell characteristics *in vivo* (*38*) *and in vitro* (*8, 29*). Based on different characterization metrices, this cell type has been referred to as mesenchymal stem cells (*8, 29, 39*), pericytes (*34*), perivascular cells (*40*), or smooth muscle cells (*26*), each of which is represented by a different xCell signature. Our evaluation using both single cell data and artificial mixtures discovered the lack of specificity of xCell’s signatures (MSC, pericyte, and smooth muscle cell) to eMSC, especially with low eMSC abundance (**Fig. 2C**), due to concurrent expression of these signatures in eSF. This observation confirms the close relationship between these two endometrial cell types and their common association with progenitor MSC and pericytes. eMSCs are implicated in endometriosis (*8, 29*) and future studies should use unique markers identified for this cell type, such as *RGS5*, *GUCY1A2*, and *NOTCH3* (*8, 26, 34*).

xCell’s fibroblast signature did not pass the permutation test despite receiving moderate ratioNext score and “onTarget” classification (**Fig. 2B**). This discrepancy may be due to many factors. Firstly, expression levels of fibroblast signature genes that passed the thresholds for ratioNext calculations (i.e. adjusted p-value and log2(fold change), **Method**) often showed only the borderline fold changes, and as Subramanian et al. explain, signatures of this nature can be expected to score poorly in a gene set enrichment based method (*41*). Furthermore, seemingly unrelated signatures such as chondrocytes, astrocytes, smooth muscle, and MSC showed highest enrichment for the eSF cluster of the single-cell data (Fig. 2B) and artificial mixture analysis confirmed that one of these cognate cell types, eMSC, could contribute to Fibroblasts-signature enrichment scores (Fig. 2C,D). Our signature evaluation method (**Fig. 1A**) thus considered output of this signature with low confidence. It’s known that eSF have their own unique phenotype that is distinct from other fibroblasts of the body in many ways, and that expression profile as well with hormonal changes in the endometrium (*8, 26, 34*). Future studies should use published dataset (*8, 26, 34*) to identify signatures that are specific to human eSF.

### Contributions of Immune Cells to Endometriosis

Our scRNAseq analysis identified 13 transcriptomically distinct immune type/subtypes in healthy endometrium, which were previously broadly categorized into lymphocytes and macrophages (*26*). The use of a signature-based method (*30*) and classical cell type markers allowed us to confidently annotate pDC and monocytes, which have not been confidently identified at single cell resolution or functionally examined in endometrium. With confirmed applicability of xCell’s signature for both cell types in endometrium, our deconvolution results revealed relative increases in pDC and monocytes during MSE in women with endometriosis (**Figs. 4 & 5**), suggesting likely involvement in inflammation. pDC have known involvement in the inflammatory response normally and in pathologic settings, through interaction with vasculature and T cells (*42*). Increased monocytes are congruent with increased expression of monocyte chemoattractant protein-1 (*MCP-1*) in endometrium of women with endometriosis (*43, 44*). Moreover, we observed a greater increase in monocytes in women with stage III-IV endometriosis, which may contribute to lower implantation rate and live birth rates compared to women with stage I-II disease (*45*).

We have previously shown that endometrial macrophages (M1 and M2) in endometriosis are predominantly pro-inflammatory (*9*). Phase- or disease-stratified abundance quantification of endometrial myeloid cell types, including macrophages, monocytes, and dendritic cells, however, are limited. Here we report the unique markers that discriminate diverse endometrial myeloid cell type/subtypes for future studies.

Our scRNAseq analysis of immune cells in healthy endometrium has identified immune cell type/subtypes that are beyond definition of xCell’s 64 signatures, such as four NK cell subtypes, one CD8+ T cell subtype, one CD4 expressing T cell subtype, and a *KIT*+ lymphocyte cell type. Intriguingly, xCell’s Th1, Th2, Tgd, and CD8+ Tem signatures, were more enriched in NK cell subtypes rather than the T cell subtypes in the single cell dataset (**Figs. 5 & S7**). Therefore, xCell outputs of these signatures across the cycle and in endometriosis likely reflect changes of endometrial NK cell abundance or phenotypes more than changes in the nominal cell types. However, these results do not conclusively suggest that these cell types are not present in endometrium with or without disease, especially considering Th1 involvement in cytokine secretion and Tgd intraepithelial presence. Rather they are likely low in abundance, and their transcriptomic signals may be interfered by those of the more abundant NK cells. Lastly, although Tregs and cDC were identified in the scRNAseq dataset of healthy endometrium and demonstrated moderate enrichment of associated xCell signatures (**Fig. 3, S7A, B**), their xCell signatures did not pass our permutation test. Enriching for aforementioned cell types/subtypes with classical markers and single cell level identification is warranted in future studies.

Notable for some xCell signatures that passed our permutation analysis are the elevated abundance scores of eosinophils, neutrophils, basophils, NKT, and immune progenitors in endometrium of women with endometriosis and their absence in women without disease and in the annotations of scRNAseq dataset of healthy endometrium. Eosinophils, initiators of inflammatory responses, were enriched in all phases of the cycle, compared to control endometrium wherein they appear mainly during menses, confirmed herein and by others (*46*). Thus, eosinophils likely contribute to the pro-inflammatory phenotype observed in bulk-tissue analysis of endometrium from women with endometriosis. Our finding of neutrophils, key participants in the innate immune response to foreign pathogens and enriched in endometrium of women with endometriosis and independent of cycle phase, compared to controls, is consistent with other reports, although others found cycle-dependence of this cell population in women with versus without disease (*47, 48*). Basophils also initiate inflammatory responses and were found herein to be enriched in endometrium of women with disease in PE and were significantly increased throughout the cycle. We are unaware of other reports on this cell type in endometrium of women with endometriosis, and this finding warrants further study.

### Comparison to Prior Work

Our results generally agree with a prior deconvolution study on endometrium from women with and without endometriosis (*49*), although, fewer xCell signatures with disease-associated changes (9 in total) were identified compared to our study. Differences may be due to our adapted usage of the deconvolution method. Additionally, this study (*49*) did not design or apply metrics for statistical significance evaluation and result interpretation or develop *de novo* identification of normal endometrial immune cell signatures to enrich the xCell data interpretation in the endometrial context.

### Strengths and Limitations of this Study

There are several limitations in this study. One is the limited sample size, especially in the ESE phase. Another limitation of our current approach is its limited capacity in inferring cell type-specific phenotypic state. Although such insights can still be inferred for phenotype-specifying signatures, such as several immune cell subtypes mentioned above, for signatures without tissue-matching phenotype specifications, such as fibroblasts, such insights cannot be obtained directly from the deconvolution results. This is remarkable as the eSF changes transcriptomically across the cycle and displays marked abnormalities in endometriosis (*8, 50*) and is a key regulator of successful embryo implantation. Other cell type deconvolution tools such as Cibersortx (*14*) provide the possibility to infer cell type-specific gene expression profiles through additive combinations of input cell type signatures. Successful application of this approach requires that highly specific cell type signatures be used and that all potential cell type signatures be included (*51*).

Further studies leveraging single cell technologies as well as integrating different types of omics measurements including proteomics, epigenetics and others will enable further corroboration of our findings and linking transcriptional phenotypes with endometriosis-associated cell types. Functional studies will help elucidate the roles these cell types play in disease.

Through integrated whole-tissue deconvolution and single cell analysis, we identified endometrial cellular compositions that are dynamic across menstrual cycle phases and altered in women with endometriosis. Guided by our signature evaluation metrics, we report cell type candidates - immune cell type/subtypes of myeloid lineage, as well as non-immune cells, including epithelial and endothelial cell types - that most likely contribute to the pro-inflammatory endometrial phenotype previously observed in women with endometriosis (*4, 7*). Our results can help guide the selection of cell types for functional evaluation of cellular mechanisms that contribute to or result from endometriosis. Moreover, our analytical framework can be used in studies of other tissue types.

## Methods

An overview of all methods is shown in **Fig. 1**.

### Experimental Data

#### Whole Tissue Microarray

Microarray data for this study were obtained from GSE51981 (4) and all analysis was carried out in R. Sample metadata for disease severity was used to classify all samples into groups of stages I-II and stages III-IV, with ambiguously mapping samples (n=1) being subsequently removed in further analyses. Sample metadata for pathology was used to classify samples as endometriosis, no pathology (which included labels NUP (no uterine pathology) and NUPP (no uterine or pelvic pathology)), or “other”, with all “other” samples being left out, as such samples represent imperfect controls, for subsequent analyses. Additional samples were removed which had ambiguous lab source annotation (n=1) or cycle-phase annotation outside of proliferative endometrium (PE), early secretory endometrium (ESE), or mid-secretory endometrium (MSE) (n=1). In the end, this led to 105 samples total, 71 from women with endometriosis and 34 from women with no uterine or pelvic pathology (controls) (**Table 1**).

#### Sorted Cell Microarray

Microarray data from purified human endometrial cell populations (stromal fibroblasts, endothelial, epithelial, and mesenchymal stem cells) isolated by fluorescence-activated cell sorting (FACS) were from previous studies: GSE73622, GSE31152, GSE48301, GSE97163 (*8, 32–34*). These were used in artificial mixes of pure cell types in determining signature specificity of the xCell signatures (see below).

#### Single Cell Transcriptomics

Endometrial single-cell RNAseq data used to evaluate xCell signatures were collected as endometrial biopsies from women without endometriosis or uterine or pelvic pathology, as previously described (*26*) (GSE111976 and SRP135922). For this study, 10x data published in (*26*) were used. Definition of endometrial cell types and subtypes is described in Extended Data Figure S1 in (*26*). Annotations of each cell with regard to participant, cycle phase, and cell type or subtype are available in a supplementary file “GSE111976_summary_10x_day_donor_ctype.csv.gz” under GSE111976.

### Microarray Normalization and Batch Correction

Background correction and quantile normalization were performed with the justRMA function of the affy package (52). Then batch correction was performed with ComBat (*53*) to reduce signals coming from the lab of origin, while protecting signals associated with disease stage and cycle phase. Direct principal components analysis (PCA) and the pvca package (54), which combines PCA with variance components analysis to estimate the proportion of variation in data that are associated with a set of potential sources, were used to assess the success of batch correction (**Fig. S1**).

### Differential Expression

Differential expression (DE) analysis was carried out via linear modeling using the Limma package (*55*) and the log-transformed and batch corrected expression matrix as input. Simple, ∼ single variable, formulas were used for linear model designs. When cutoffs for significant differential expression were used, they were FDR < 0.05, and abs log2 fold change > 1. Such analysis was run on various stratification of the data (**Figs. 1, S2 & S3**). For unstratified (all samples) analysis, DE was run 1) between disease versus control, 2) between stages I-II or stages III-IV versus control, and 3,4,5) between all combinations of pairwise cycle phase comparisons (PE vs ESE, PE vs MSE, ESE vs MSE). DE was also run between stages I-II versus stages III-IV, but zero genes met DE cutoffs. For stage-stratified analysis, DE was run separately on control samples only, stages I-II samples only, and stages III-IV samples only between all combinations of pairwise cycle phase comparisons. Lastly, for phase-stratified analysis, DE was run separately for PE samples only, ESE samples only, or MSE samples only, 1) between disease versus control, and 2) between stages I-II or stages III-IV versus control.

### Gene Pathway Enrichment Analysis

Pathway enrichment analysis was performed on the Broad’s hallmark gene sets (obtained via https://www.gsea-msigdb.org/gsea/msigdb/collections.jsp#H) by gene set enrichment analysis. This was carried out with the fgsea function of the fgsea package (*56*) on log2 fold changes of all genes (both significant and non-significant), for all stratifications and differential expression comparisons, with additional parameters: minSize = 15, eps = 0, and maxSize = 1500. Pathways with FDR corrected p-values below 0.05 were considered differentially enriched (**Fig. S4**).

### Cell Type Enrichment

Of the numerous deconvolution and enrichment methods, those that attempt to deconvolve a sample into additive mixtures of reference cell type signatures have a strong reliance on both concordance between reference signatures and cell type profiles of the target tissue, as well as on the presence of reference signatures for all cell types which might exist in the target sample. Given that we could not be certain that we would include signatures for all cell types which might exist in the endometrium, and that many immune cells profiled from the endometrium have shown non-canonical transcriptional profiles, we chose to use xCell’s enrichment-based approach which is more robust to signature absence and inconsistencies, and includes signatures for 64 human immune and stromal cell types (including adaptive and innate immune cells, hematopoietic progenitors, epithelial cells, and extracellular matrix cells derived from thousands of expression profiles) (*25*). xCell was run on the log-transformed and batch-corrected expression data of human endometrial tissue described above. Due to uncertainty in the applicability of xCell signatures to endometrial tissue, only the rawEnrichmentAnalysis and transformScores steps were utilized for calculation of enrichment scores. The spillOver adjustment step was not utilized due to notable deviations between xCell signatures versus nominal endometrial cell profiles which would have been carried over into the compensation matrix derived from in silico mixtures of reference cells.

### Filtration of cell type signatures based on permutation analysis

As discussed by its authors, xCell often produces non-zero scores, which may result in false-positive interpretation for non-existent cell types or unsuitable signatures. xCell signatures that might not apply well to endometrial samples were first identified based on comparison to a permuted background distribution (**Figs. S5 & S6**). A background distribution of enrichment scores was generated for every cell type signature, and for each cycle phase, by running xCell with 1000 permutations of our expression matrix where rownames (gene symbols) were shuffled. Significance testing was then performed for each cycle phase among control or disease samples individually. Median enrichment scores of all iterations, among the current stratification samples, formed the background distribution for each cell type. For a given stratification, a cell type signature was then considered as expressed if the “true” median enrichment score, from the non-permuted data, was greater than the 90th quantile of its background distribution. To ensure that cells present in only certain conditions might still be accurately assessed, this filtering procedure was run on a per-disease status and per-phase basis, and cell type signatures were retained for future analyses as long as the median enrichment score was above the background cutoff for at least one stratification.

### Evaluation of xCell signatures using single cell measurements of endometrial tissue from women without endometriosis

Meanwhile, even when an xCell score is statistically significant, its contributing xCell cell type signature may not be specific to its nominal cell type target in the tissue of interest, due to inter-tissue variability of the same cell type or ambiguity in cell type naming.

To test sensitivity, for each of the 64 xCell signatures, a signature score was calculated with respect to each endometrial cell type identified in the scRNAseq dataset (*26*). To identify differentially abundant cells, Wilcoxon’s rank sum test (two.sided) was performed, and fold change (FC, dummy variable = 10^-2) was calculated between cells from an endometrial cell type and the remaining cells. P-values obtained from Wilcoxon’s rank-sum test were adjusted for multiple comparison by the Benjamini– Hochberg’s procedure to obtain p.adj. A signature score was quantified as the percentage of genes in the given xCell signature that were differentially expressed between cells in an endometrial cell type compared to the remaining cells (p.adj < 0.05, log2(FC) > 1). For each of the xCell signature, the resulting score was normalized by the median of scores of all eight endometrial cell types identified in the single cell dataset (Normalized s_i, j_ = s_i, j_ / Med(s_i_,_1_, s_i_,_2_, … s_i_,_8_), where i is an xCell signature and j an endometrial cell type identified in the single cell dataset).

Each xCell signature was categorized as either “reference” or “no-reference”, based on whether there is an endometrial cell type or subtype in the single cell dataset that the signature is potentially targeting. A map between each xCell signature and each endometrial cell type was constructed to describe this relationship (**Fig. 2B, boxes**). As shown in **Fig. 1**, we kept this relationship relatively broad such that a signature is considered targeting a single cell type/subtype if it targets directly the identified endometrial cell type, or a sub-category of the identified cell type, or a related category of the identified cell type, to account for ambiguity in naming cell types and for potential existence of subtypes within the annotated cell populations.

Two specificity score metrics were then established. Given the target map, for the first specificity metric, “onTarget”, an xCell signature was tagged as “on-target” if the highest-ranking endometrial cell type from the single cell expression data matches the cell type targeted by the xCell signature and “off-target” otherwise. Signatures without a clear reference cell type within the single cell dataset were given an ‘NA’ label (**Fig. 2B**).

Separately, to evaluate how specific an xCell signature is to the endometrial cell type it represents, we calculated a “ratioNext” score representing the ratio between the highest and the 2nd highest ranking signature scores. Importantly, to avoid over-penalizing, if subtypes exist for the highest-ranking cell type (e.g., epithelial cells), scores in the subtypes were ignored in determining the 2nd highest signature score (**Fig. 2B**).

### Identification and annotation of 13 immune cell type/subtypes from healthy human endometrium

Dimensional reduction was performed on cells from the two clusters annotated as “Lymphocytes” and “Macrophages” in the original analysis (*26*) using Seurat’s (v3.2.0)(*57*) implementation of uniform manifold approximation and projection (UMAP). Specifically, top 2000 variable genes among the immune cells were identified via FindVariableFeatures(). Principal component analysis was performed via RunPCA() on the top variable genes. Dimension reduction was performed on the top 20 principal components (PCs) via RunUMAP() based on the distribution of variances explained by the top PCs. Cell type/subtypes were identified using Seurat’s FindNeighbors(dims = 1:20) and FindClusters(resolution = 0.6). For each identified cell type/subtype, FindNeighbors() and FindClusters() were iterated one additional round to identify further heterogeneity. A cluster is classified as a candidate immune cell type/subtype if it can be defined by statistically significant uniquely expressing markers. pDC and B cells were present in 4 samples, macrophages were present in 7 women and the rest of identified immune type/subtyes were in all 10 women.

For each identified cell type/subtype, uniquely expressing genes were found via FindAllMarkers(only.pos = TRUE, min.pct = 0.25, logfc.threshold = 0.25, test.use = “wilcox”, slot = “data”) and ordered based on log2FC.

As elaborated in the text, annotation of each immune cell type/subtype was performed through iterative evaluation of classical marker expression, signature level scoring of xCell’s immune cell signatures, and RNA expression pattern of uniquely expressing genes identified above reported by the Human Protein Atlas (*31*). For signature scoring, we used the method reported in (*30*). Briefly, for each xCell’s immune signature, the score was quantified as the ratio between transcripts (UMI) that encode genes in the signature to all transcripts (UMI) detected in each single cell. We further examined the distribution of each signature in each identified immune cell type/subtype.

### Validation of xCell approach using artificial mixtures from sorted cells

Microarray expression data from four cell types of FACS-purified endometrial cells (from participants with and without endometriosis) were used to generate 20 different artificial mixtures with varying proportions of each cell type. The microarray expression data from endothelial cells (n=11 samples), epithelial cells (n=7), mesenchymal stem cells (n=28), and stromal fibroblasts (n=31) were first summarized by their median expression for all probes. These median cell profiles were then additively combined into 20 different mixtures in which one cell type made up 10, 30, 50, 70, or 90% of the mixture, and the remaining cell types made up the remaining 90, 70, 50, 30, or 10%, respectively. xCell was then run on these mixtures both with and without the spillOver step (**Fig. 2C, D**).

### Differential cell type enrichment analysis

Log2 enrichment ratios (log2ER), between groups, were calculated for each cell type signature. P-values were generated by performing two-sided Mann-Whitney U tests between enrichment scores of all samples, between groups. These were then corrected for multiple hypothesis testing via the FDR method based on the number of signatures assessed. FDR < 0.05 was the sole cutoff used for differential cell type enrichment. Such analysis was run on the same stratifications of the samples and for the same comparisons for each of those stratifications, as for differential gene expression analysis, described previously (**Fig. 1**).

## Supplemental Tables

**Table S1: Differentially Expressed Genes, Case/Stage vs Control**

Excel file with tabs: Unstratified DvC, PE sample DvC, ESE sample DvC, MSE sample DvC, Stages I-II vs control, Stages III-IV vs control

**Table S2: Pathway Analysis, Case/Stage vs Control**

Excel file with tabs: Unstratified DvC, PE sample DvC, ESE sample DvC, MSE sample DvC, Stages I-II vs control, Stages III-IV vs control

**Table S3: Differentially Expressed Genes, Between Phases**

Excel file with tabs: PE vs. ESE (Controls), PE vs. MSE (Controls), MSE vs. ESE (Controls), PE vs. ESE (Cases), PE vs. MSE (Cases), MSE vs. ESE (Cases)

## Supplemental Figures

**Figure S1:**
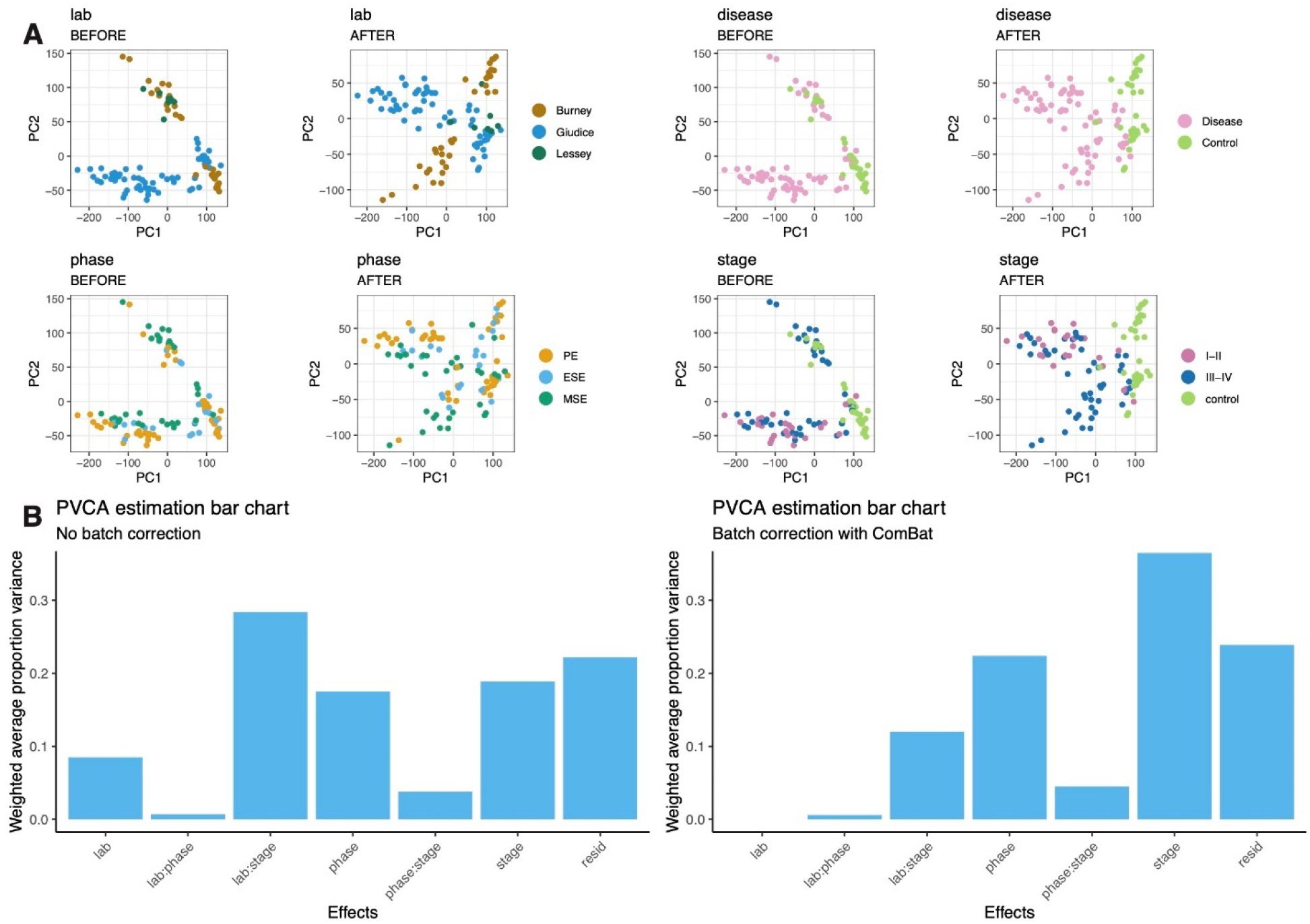
Batch correction reduces contributions from samples’ lab origin. **A.** Principal component Analysis (PCA) of gene expression data (left) before and (right) after batch correction with ComBat with samples colored by different metadata. **B.** Bar plots showing the estimated variance associated with technical variables (“lab” of origin), biological variables (phase & stage), or interaction terms representing combinatorial contributions of these variables, (left) before and (right) after batch correction with ComBat.

**Figure S2.**
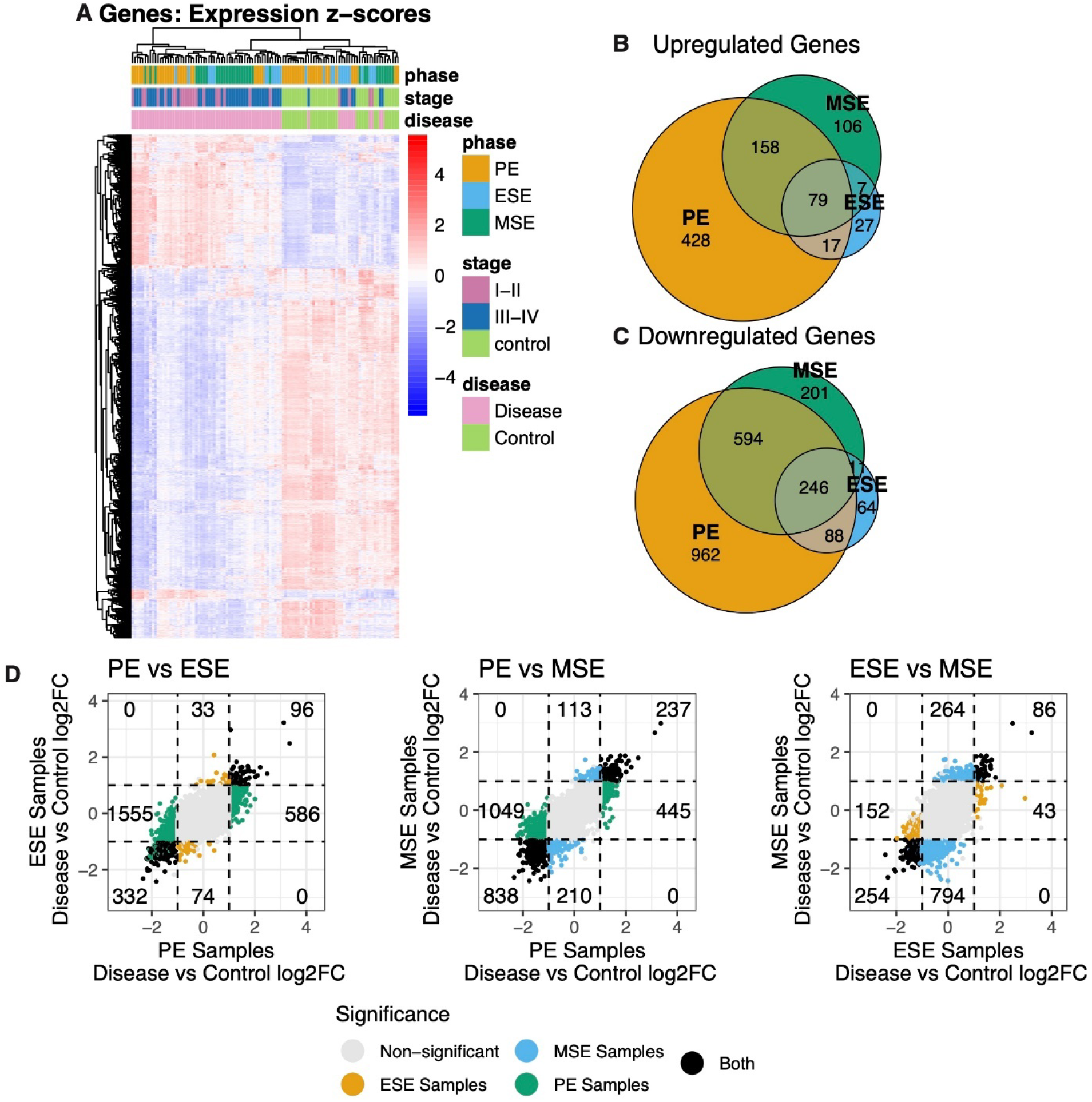
Disease versus Control Comparison on the Gene Level. Differential gene expression was performed between disease versus control and Stages I-II or Stages III-IV versus control within various stratifications of the samples. **A.** Heatmap showing, for all samples, (A) relative, z-score, expression of all genes determined to be differentially expressed**. B, C.** Venn diagrams comparing the composition of genes (B) up- or (C) down-regulated in disease versus control samples of each phase. **D.** Fold-fold plots comparing the log2 fold changes between disease versus control samples in the different phases.

**Figure S3:**
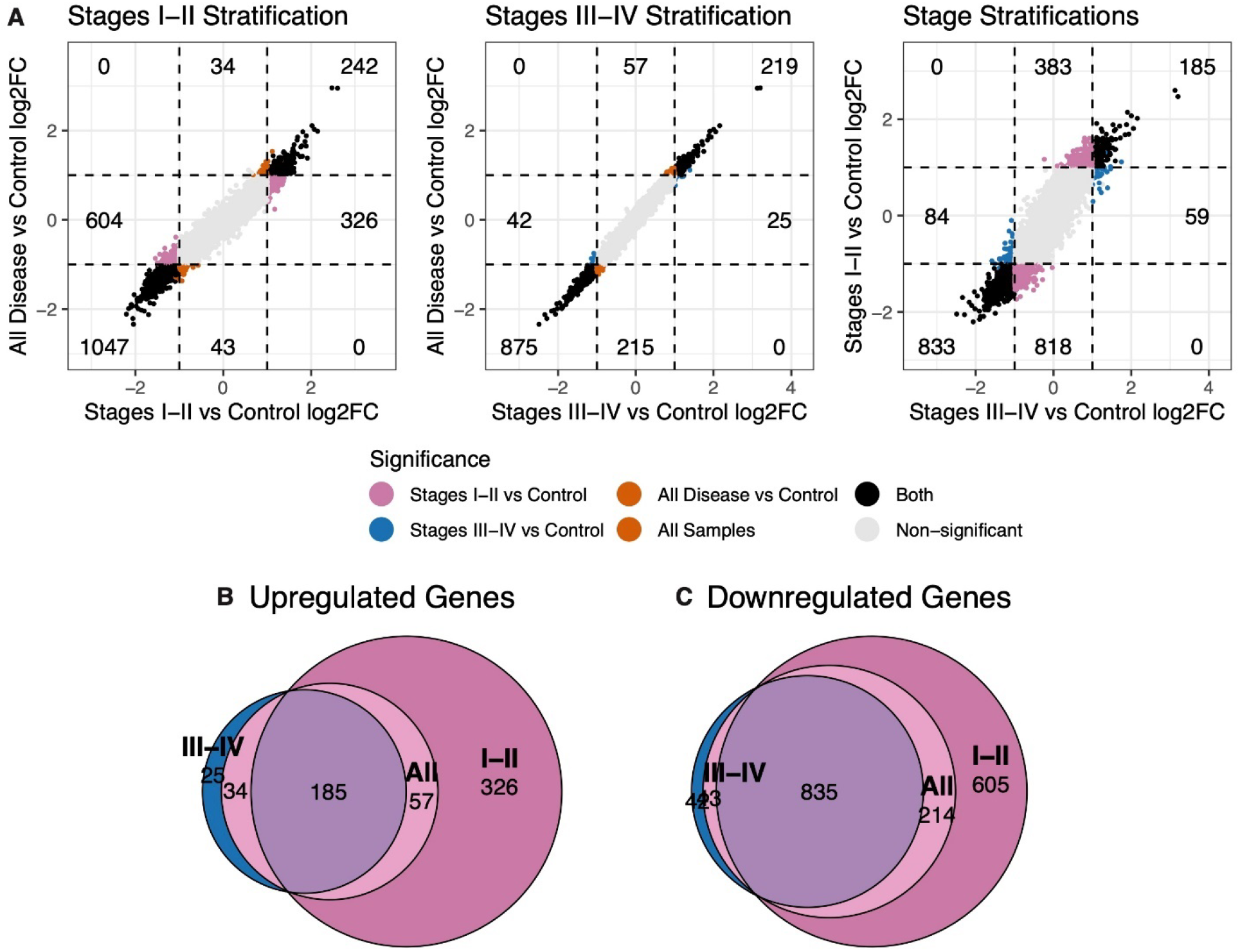
Disease and Stage versus control signature comparisons. Differential gene expression was performed between disease versus control and Stages I-II or Stages III-IV versus control samples. **A.** Fold-fold plots comparing the log2 fold changes between each of these comparisons. **B, C.** Venn diagrams comparing the composition of genes (B) up- or (C) down-regulated in disease/stage versus control samples.

**Figure S4.**
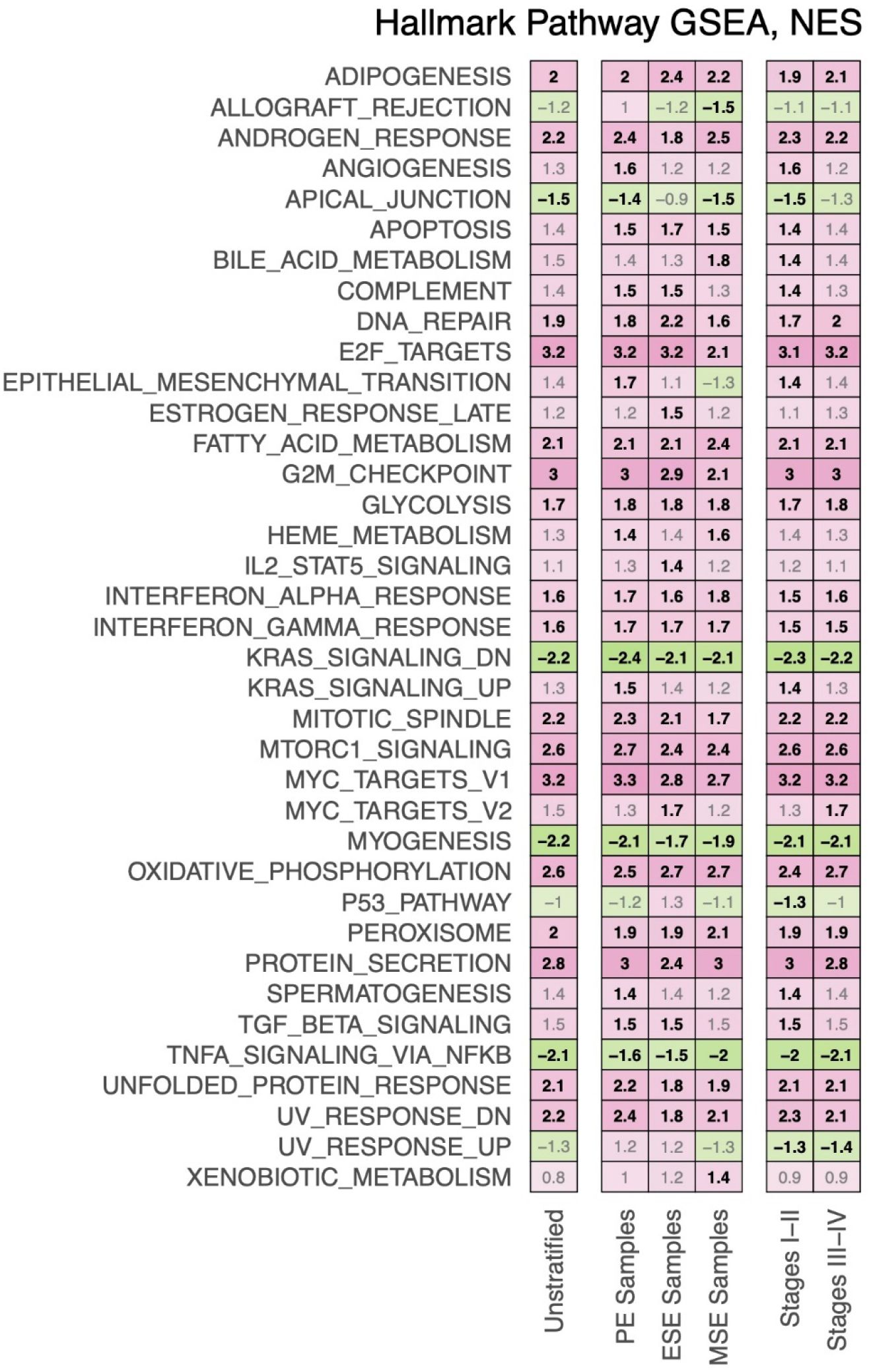
Pathway analysis was performed by GSEA using log2FC as input. Heatmap of GSEA normalized enrichment scores (NES) for hallmark pathways where only pathways with at least one significant enrichment are shown, and pathways related to the immune system are bolded. Case vs control comparisons stratified by phase and disease stage. NES with black color for statistically significant enrichments, and grey color for non-statistically significant enrichments.

**Figure S5:**
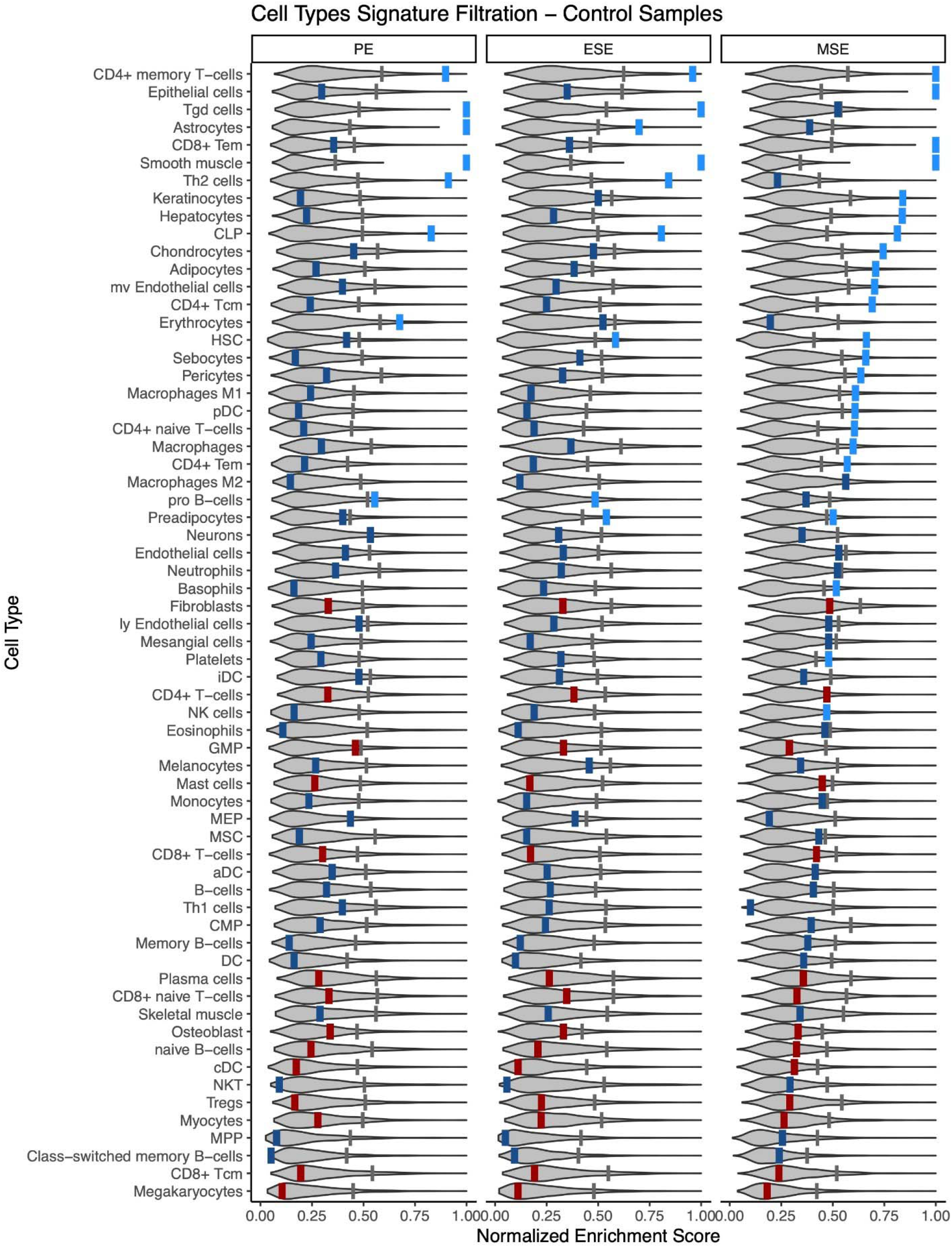
xCell Signature filtration based on a permuted background distribution (control samples) Per-phase background distributions for cell type enrichment scores (ES) were generated by permuting gene symbols (row names) of the transcriptome matrix and then running xCell, 1000 times, then taking the median ES per phase of control and disease samples, in each iteration. Shown here are violin plots of these background distributions for each cell type signature, among control samples. Thin gray vertical lines represent the 90th quantile of these background distribution values. Thicker vertical lines represent the true median ES (from non-permuted data) for the given cell type with colors: Light blue = true median ES was greater than the background cutoff for this phase (signature retained); Dark blue = true median ES was less than the background cutoff for this phase, but not for another phase (signature retained); Dark red = true median ES was less than the background cutoff for all phases (signature filtered out for subsequent analyses)

**Figure S6:**
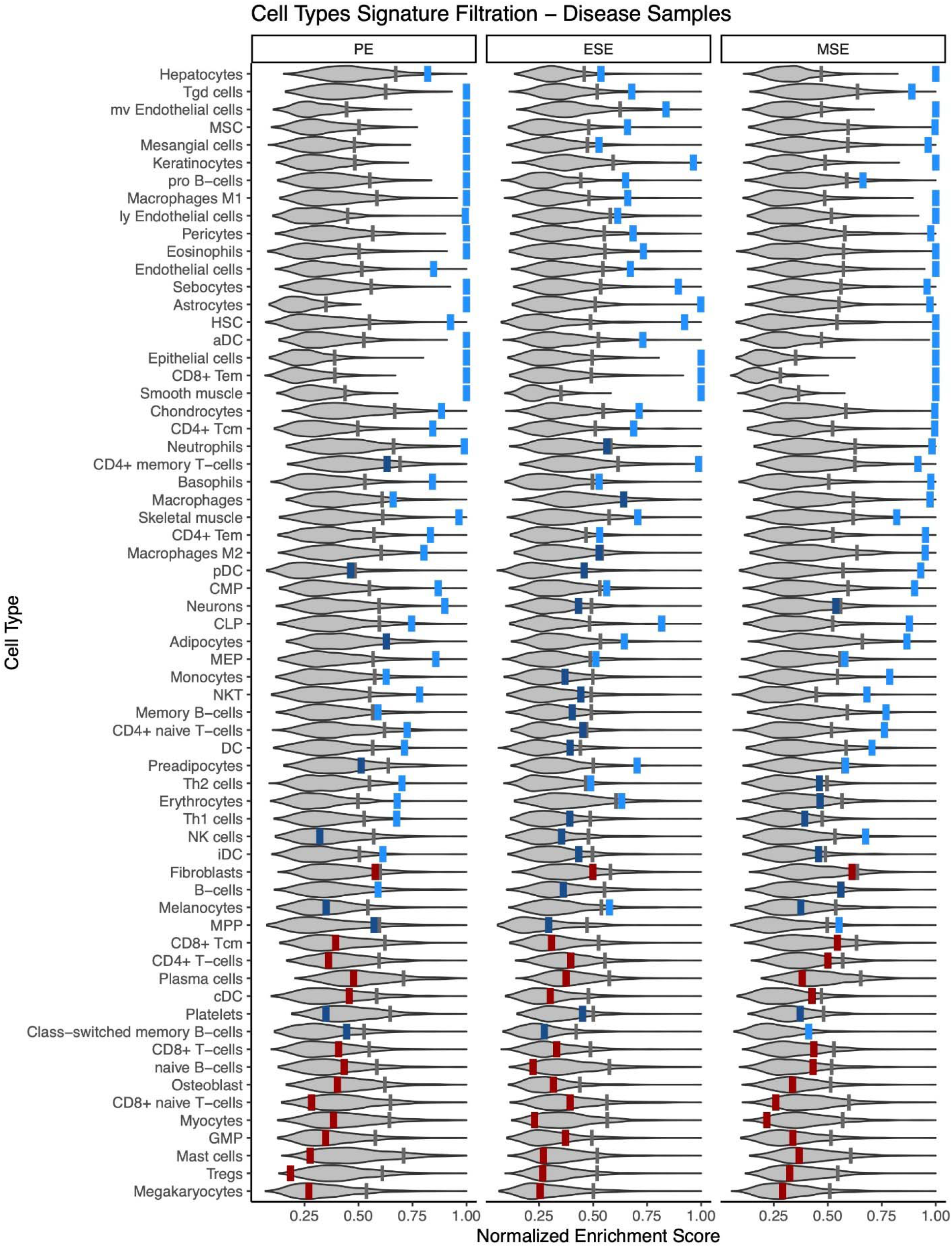
xCell Signature filtration based on a permuted background distribution (disease samples) Per-phase background distributions for cell type enrichment scores (ES) were generated by permuting gene symbols (row names) of the transcriptome matrix and then running xCell, 1000 times, then taking the median ES per phase control and disease samples, in each iteration. Shown here are violin plots of these background distributions for each cell type signature, among samples from women with endometriosis. Thin gray vertical lines represent the 90th quantile of these background distribution values. Thicker vertical lines represent the true median ES (from non-permuted data) for the given cell type with colors: Light blue = true median ES was greater than the background cutoff for this phase (signature retained); Dark blue = true median ES was less than the background cutoff for this phase, but not for another phase (signature retained); Dark red = true median ES was less than the background cutoff for all phases (signature filtered out for subsequent analyses)

**Figure S7.**
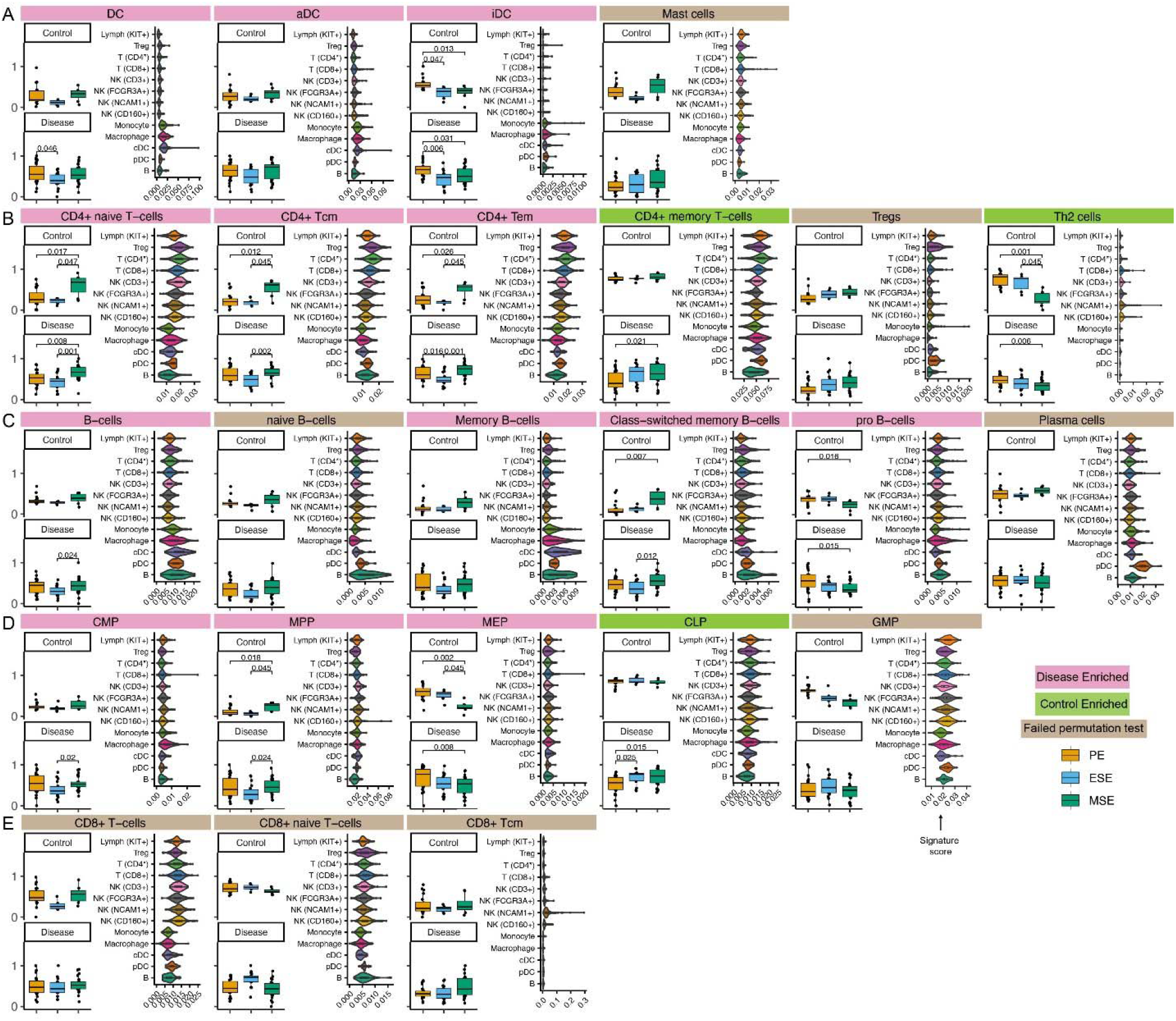
Deconvolution results and signature score distribution of xCell immune signatures of (**A**) myeloid lineage, (**B**) T cell types (**C**) B cell types (**D**) progenitors and (**E**) CD8+ T cell types. Signatures shown in Figs. 3 and 5 are not repeated here.

**Figure S8.**
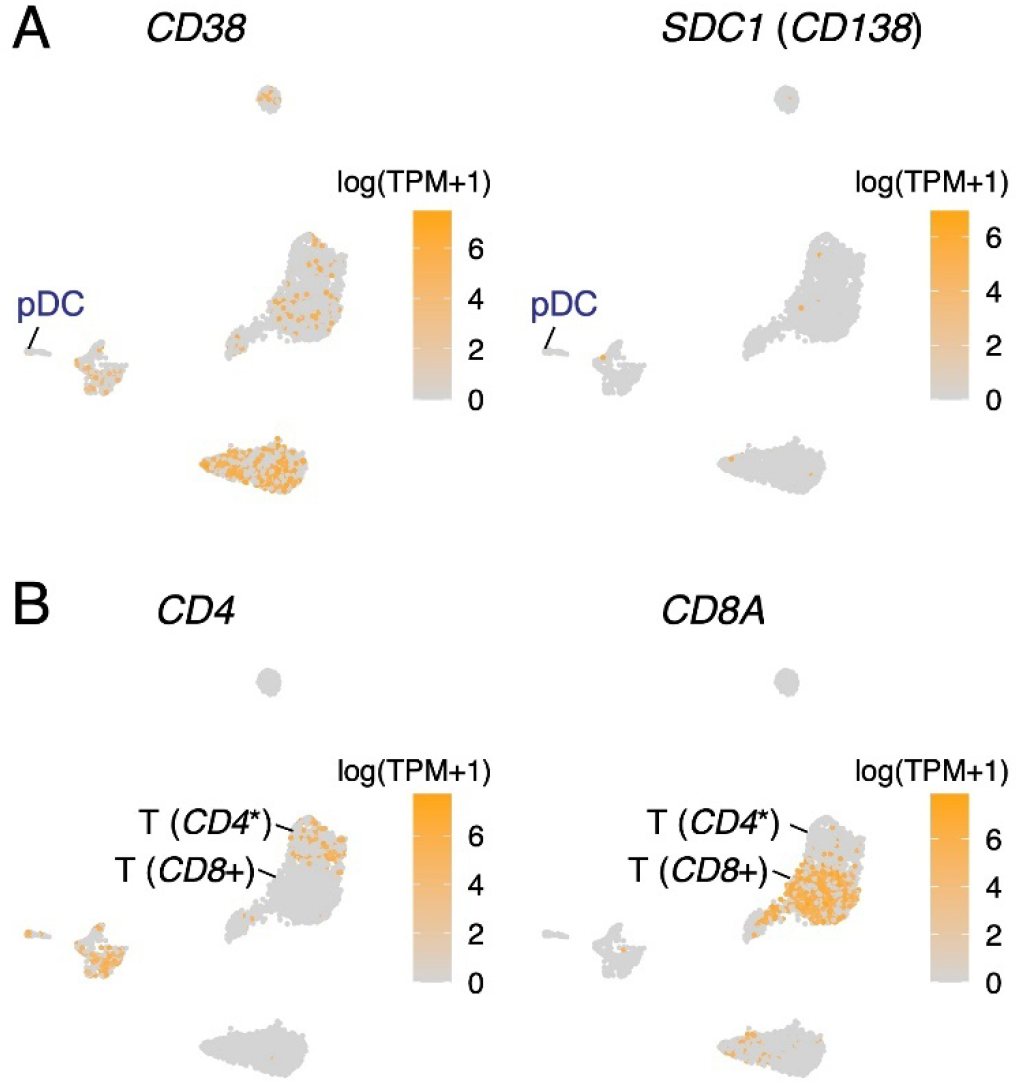
Absence of expression of classical plasma cell markers in pDC (A) and expression of *CD4* and *CD8A* in CD4* T and *CD8*+ T cells, respectively (B), identified in healthy human endometrium. TPM: Transcript per million. Log is in natural log. *CD4*\*: *CD4* was uniquely but sparsely expressed in the cell subtype (**B**) and hence was not identified as a top uniquely expressed gene in (Fig. 3B).

**Figure S9.**
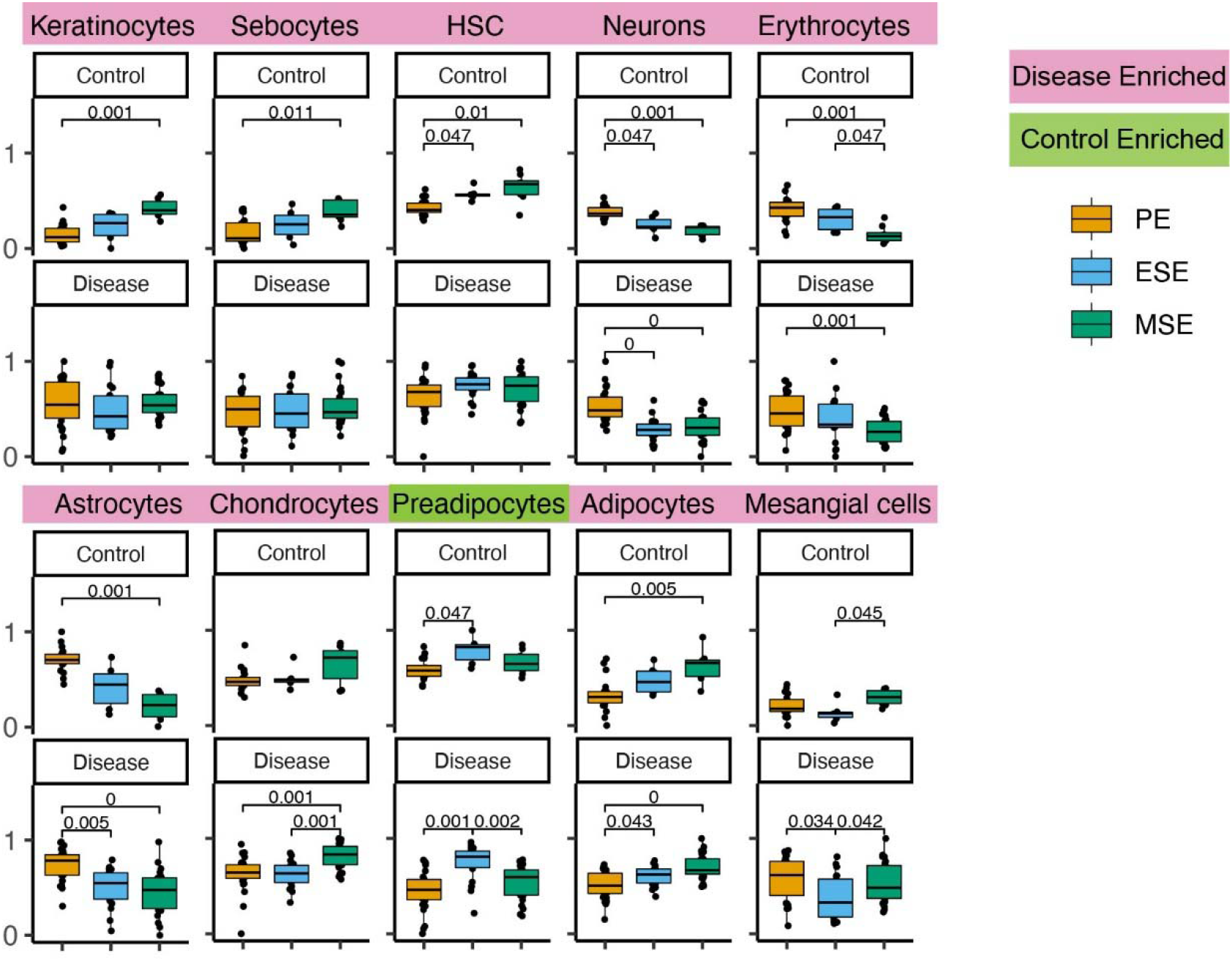
Deconvolution results of select xCell signatures. Enrichment scores are on the y-axis. Signatures shown in **Figs. 3** and **5** are not repeated here.

## Acknowledgements

The work has been supported in part by National Institutes of Health, *The Eunice Kennedy Shriver* National Institute for Child Health and Human Development, National Centers for Translational Research in Reproduction and Infertility P50 HD055764. We would like to thank Sally Mortlock, Grant Montgomery, Gabriela Fragiadakis and Dvir Aran for useful discussion.

