## Supplementary figures and images for "Whole-tissue deconvolution and scRNAseq analysis identify altered endometrial cellular compositions and functionality associated with endometriosis"

### Supplemental Fig 1

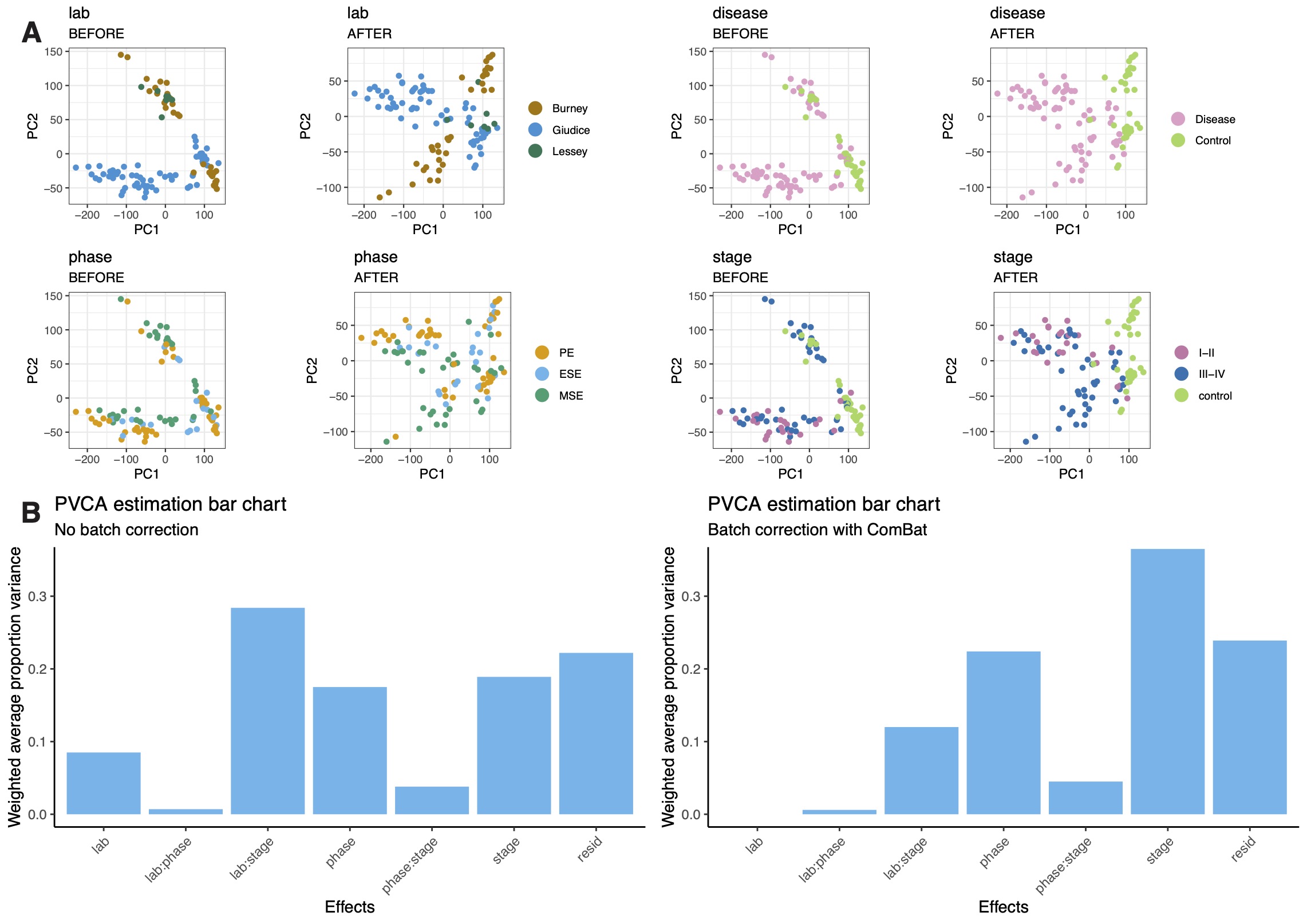

### Supplemental Fig 2

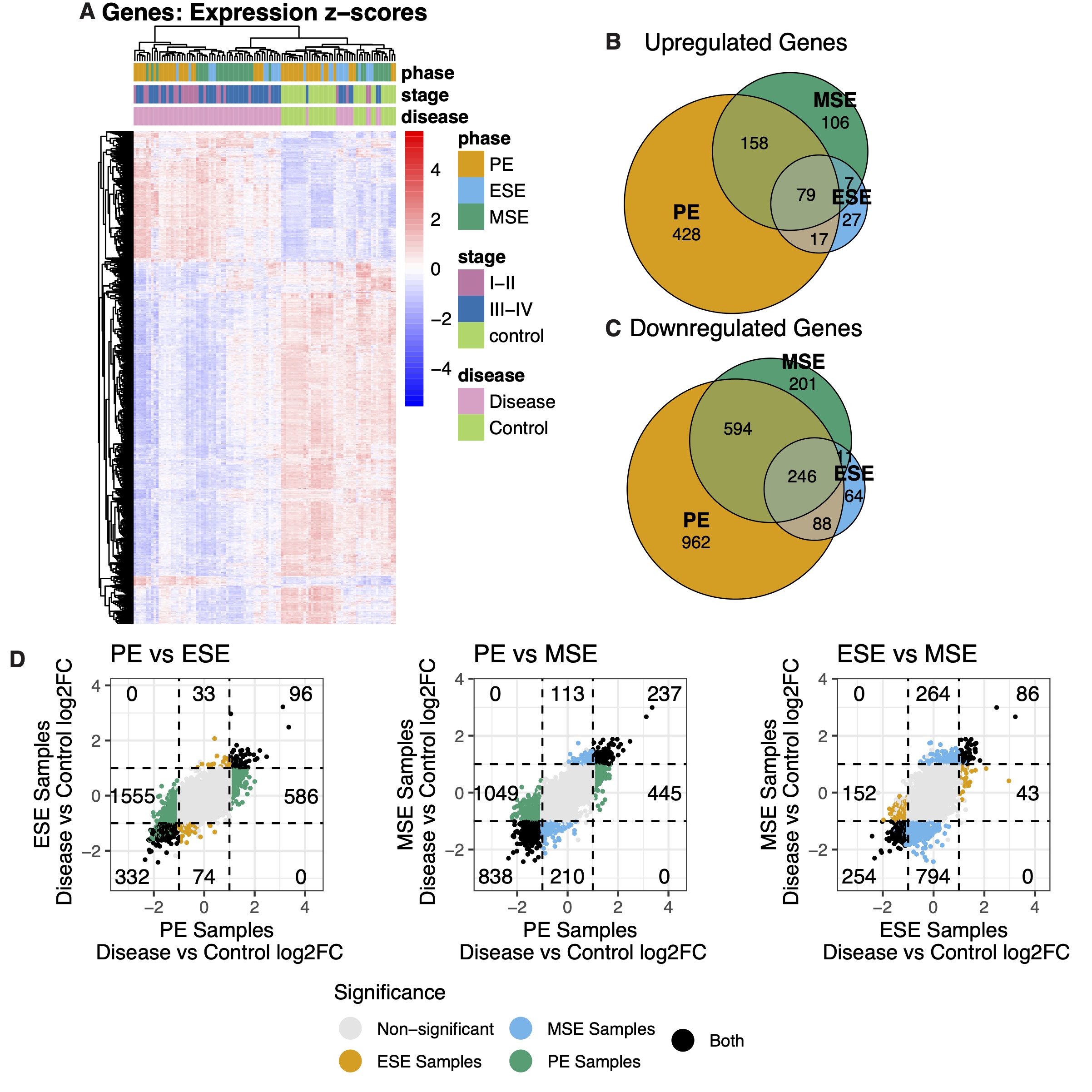

### Supplemental Fig 3

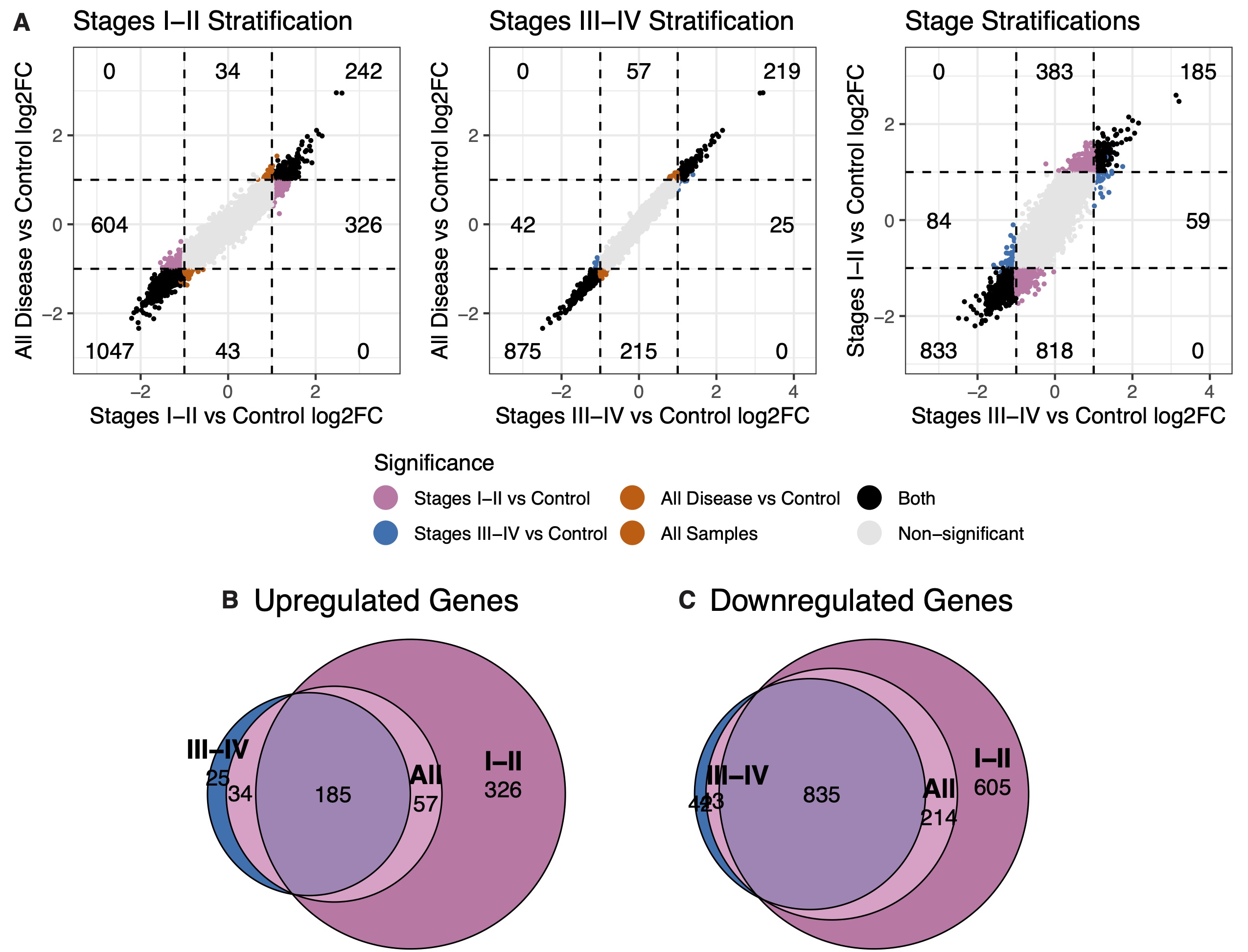

### Supplemental Fig 4

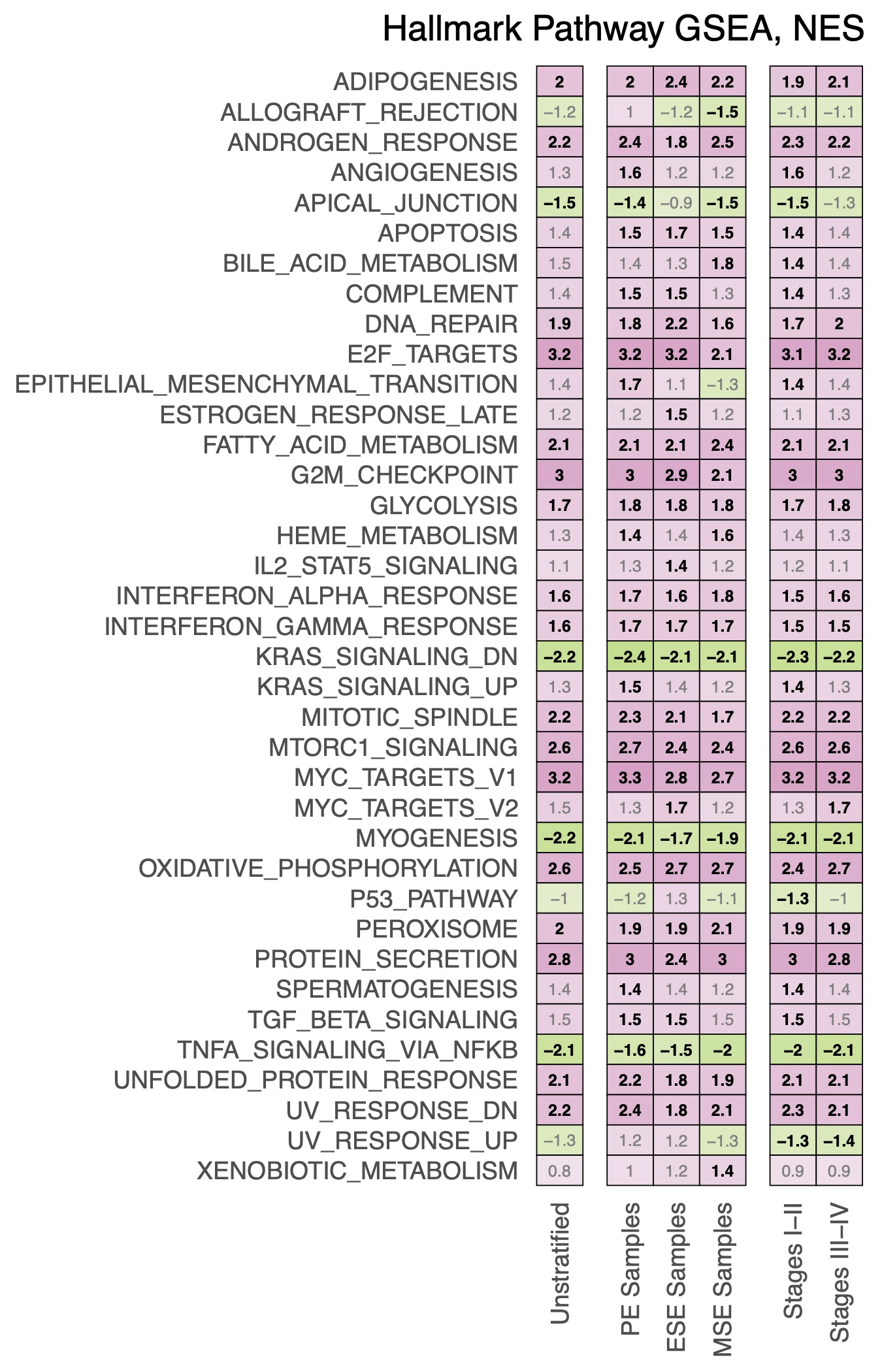

### Supplemental Fig 5

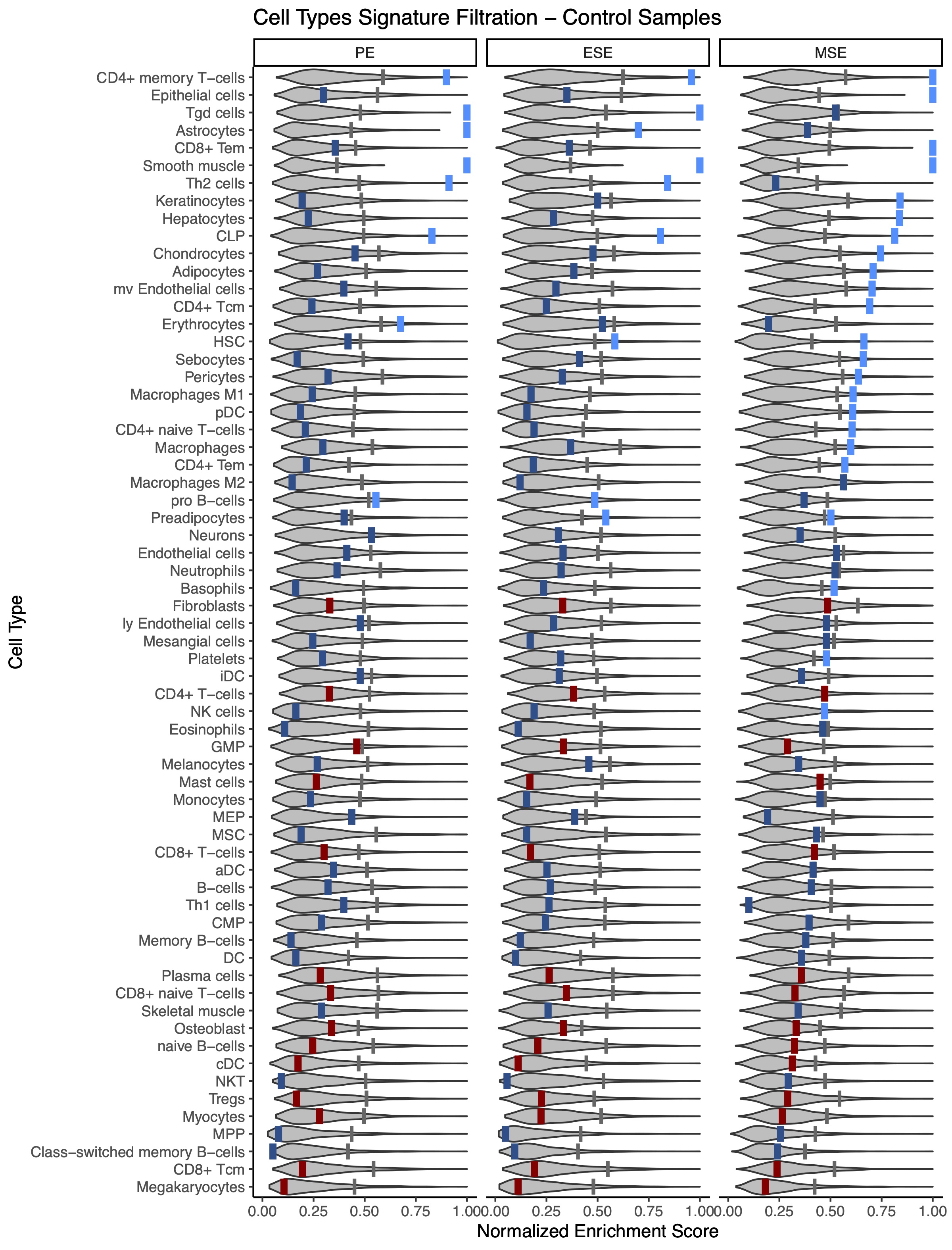

### Supplemental Fig 6

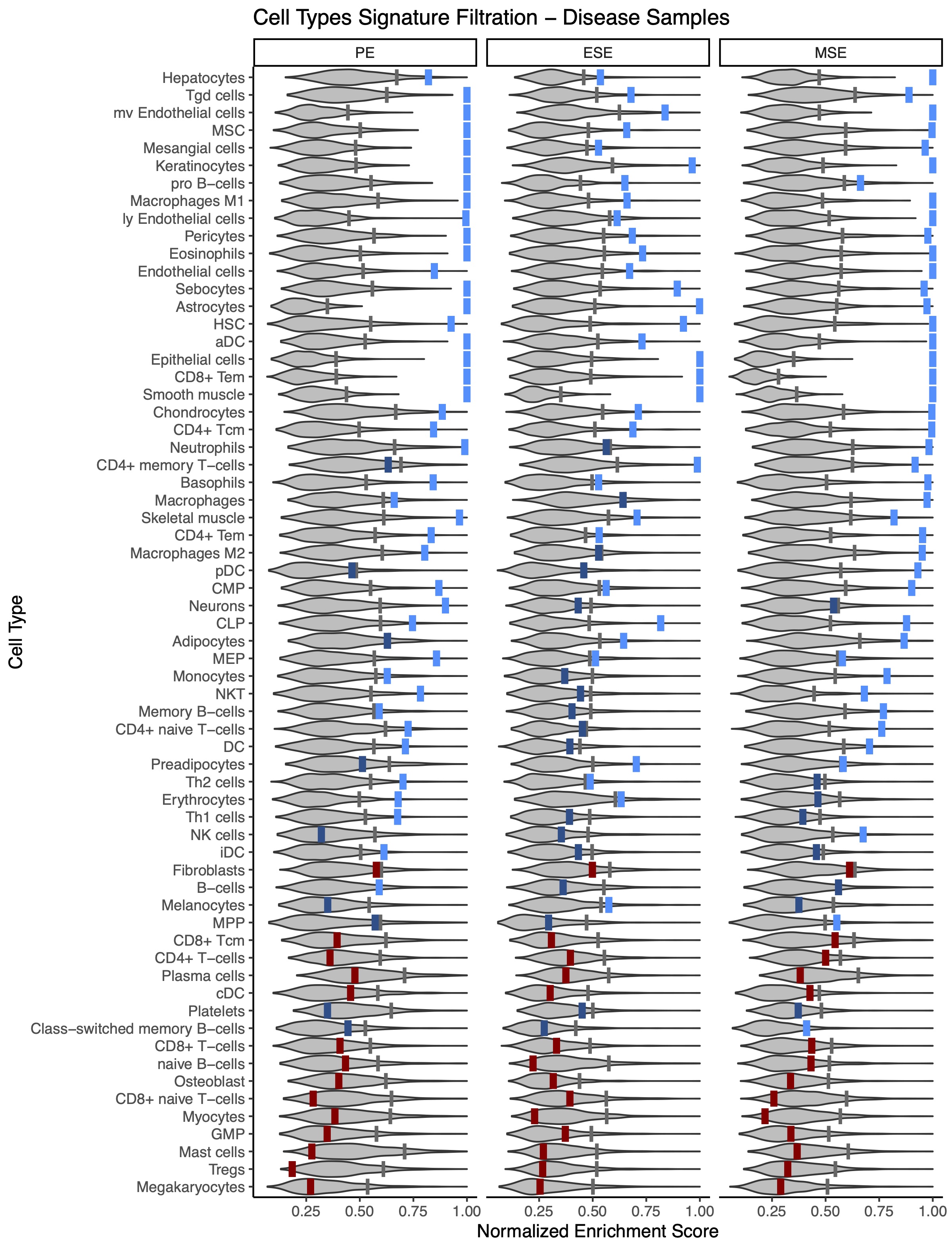

### Supplemental Fig 7

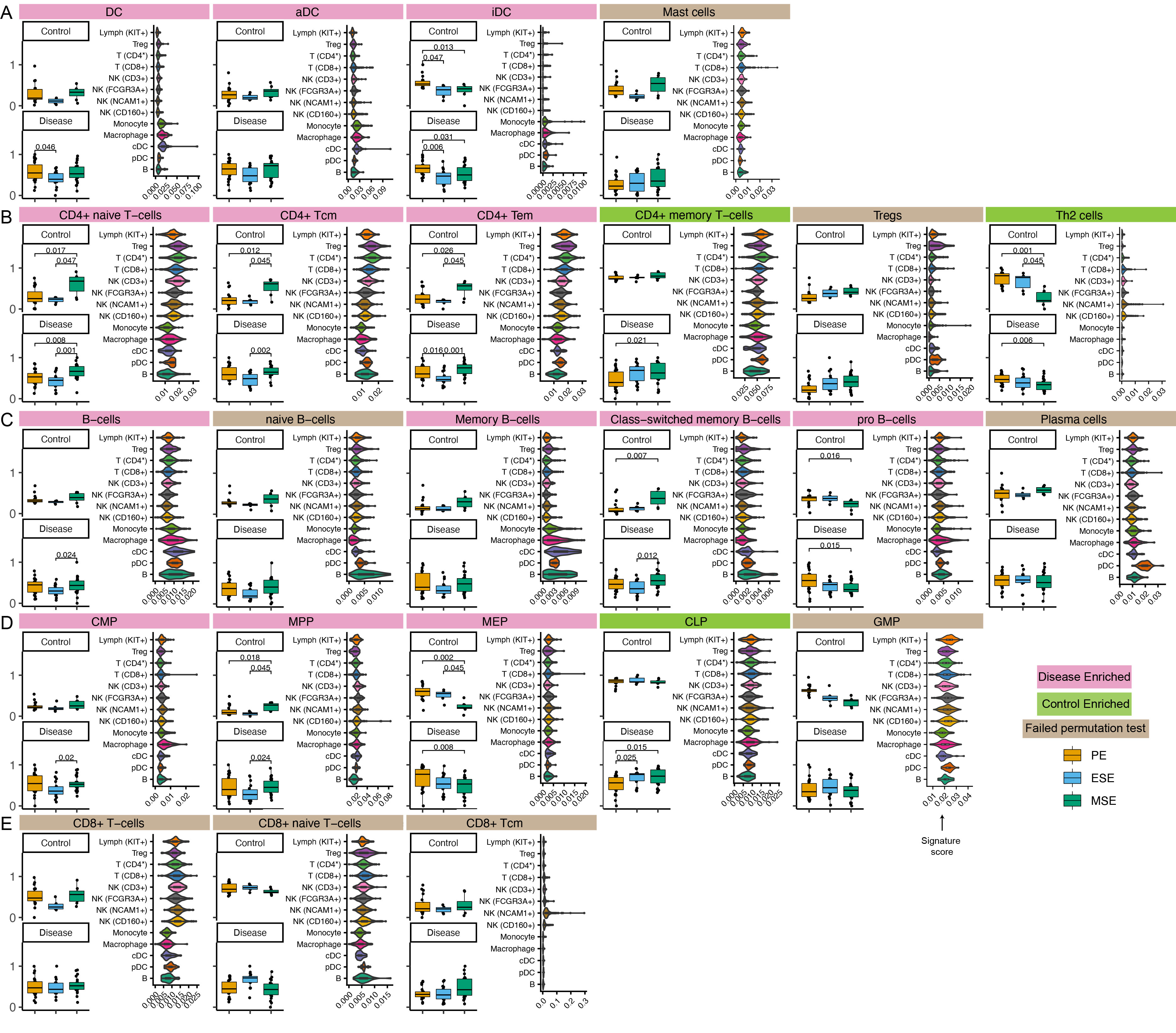

### Supplemental Fig 8

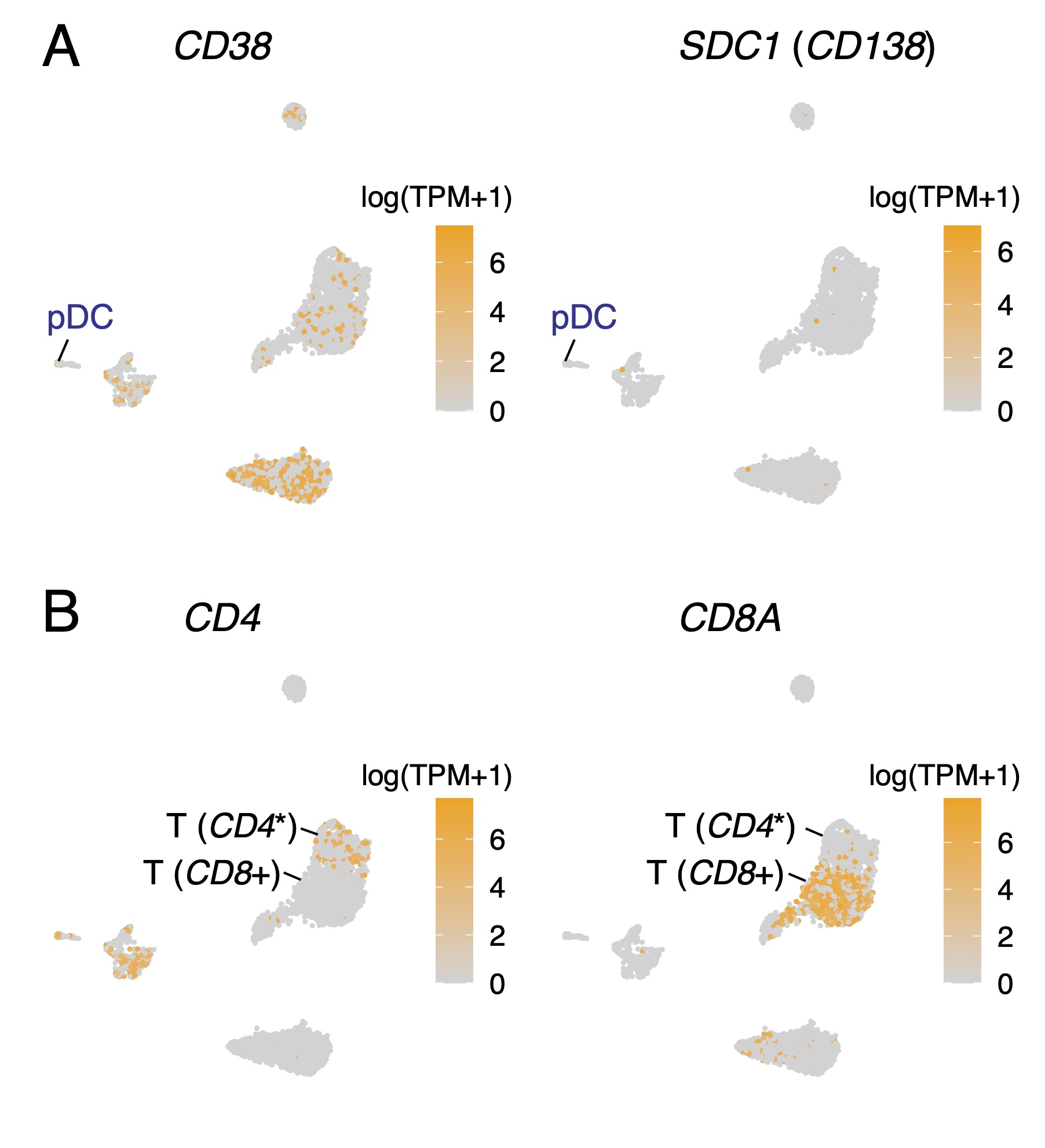

### Supplemental Fig 9

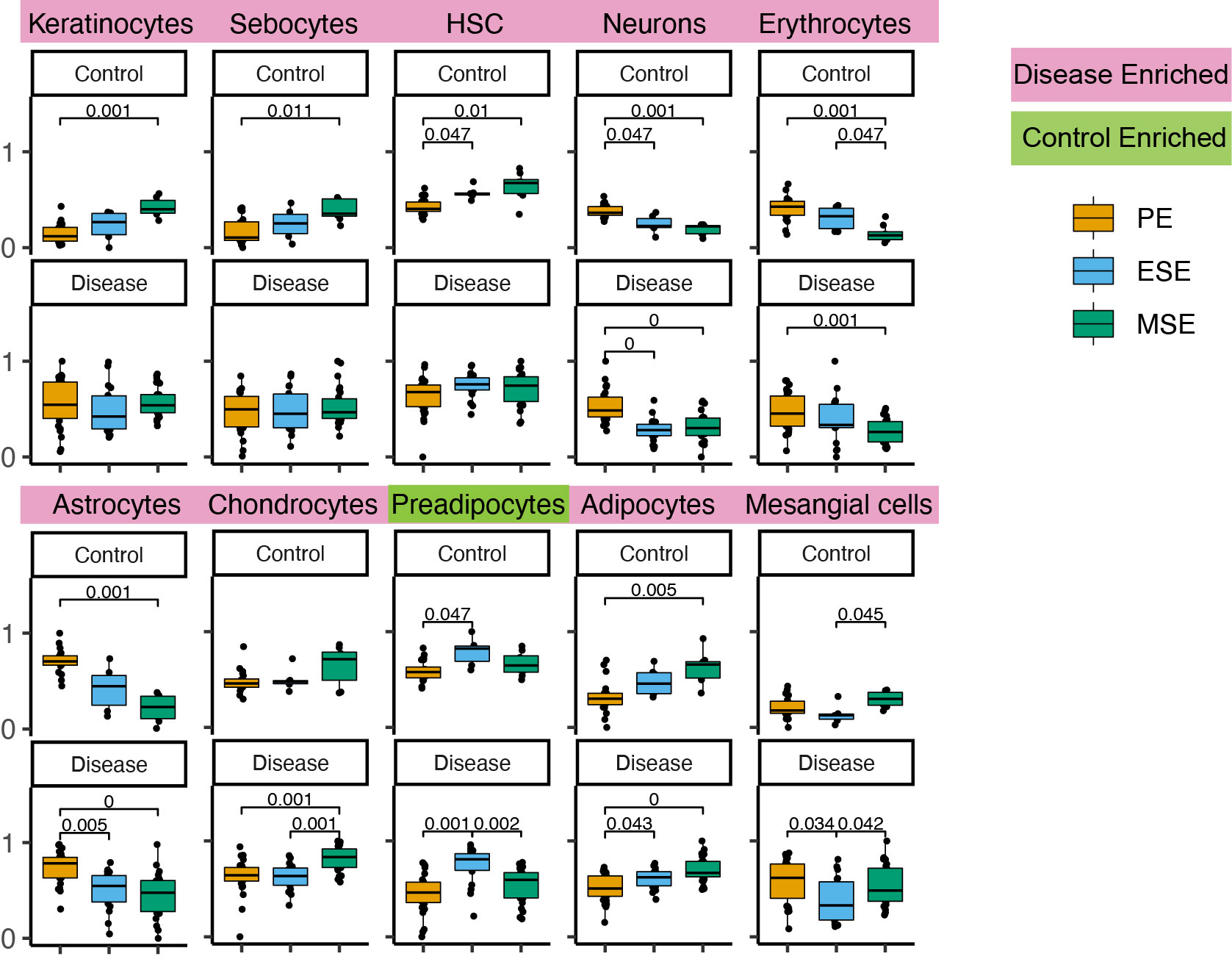
